# Conservation and Convergence of Genetic Architecture in the Adaptive Radiation of *Anolis* Lizards

**DOI:** 10.1101/2021.02.18.431064

**Authors:** Joel W. McGlothlin, Megan E. Kobiela, Helen V. Wright, Jason J. Kolbe, Jonathan B. Losos, Edmund D. Brodie

**Author notes:** The authors wish to be identified to the reviewers.

## Abstract

The **G** matrix, which quantifies the genetic architecture of traits, is often viewed as an evolutionary constraint. However, **G** can evolve in response to selection and may also be viewed as a product of adaptive evolution. Convergent evolution of **G** in similar environments would suggest that **G** evolves adaptively, but it is difficult to disentangle such effects from phylogeny. Here, we use the adaptive radiation of *Anolis* lizards to ask whether convergence of **G** accompanies the repeated evolution of habitat specialists, or ecomorphs, across the Greater Antilles. We measured **G** in seven species representing three ecomorphs (trunk-crown, trunk-ground, and grass-bush). We found that the overall structure of **G** does not converge. Instead, the structure of **G** is well conserved and displays a phylogenetic signal consistent with Brownian motion. However, several elements of **G** showed signatures of convergence, indicating that some aspects of genetic architecture have been shaped by selection. Most notably, genetic correlations between limb traits and body traits were weaker in long-legged trunk-ground species, suggesting effects of recurrent selection on limb length. Our results demonstrate that common selection pressures may have subtle but consistent effects on the evolution of **G**, even as its overall structure remains conserved.

## Introduction

Genetic variation translates natural selection into evolutionary change (Falconer and MacKay 1996; Roff 1997). Understanding the nature of such genetic variation and the processes that shape it has been a goal of evolutionary biology since the early days of population genetics (Dobzhansky 1937). In evolutionary quantitative genetics, the pattern of genetic variation in a population is described by the genetic variance-covariance matrix **G**, which predicts the multivariate response to phenotypic selection (Lande 1979). The **G** matrix can be used to make accurate predictions of short-term evolutionary change (Grant and Grant 1995), but its utility for making long-term predictions is more suspect because **G** itself may evolve during adaptive evolution (Turelli 1988; Steppan et al. 2002). Although early theory argued for the stability of **G** (Lande 1980), more recent theoretical (Agrawal et al. 2001; Jones et al. 2003; 2004, 2012, 2014; Revell 2007; Arnold et al. 2008; Pavličev et al. 2011; Melo and Marroig 2015; Melo et al. 2016) and empirical results (Steppan et al. 2002; Cano et al. 2004; Doroszuk et al. 2008; Hine et al. 2009; Eroukhmanoff and Svensson 2011; Björklund et al. 2013; Careau et al. 2015; Penna et al. 2017) suggest that **G** can and does evolve, sometimes rapidly. Given enough time, selection is expected to align **G** with the adaptive landscape (Cheverud 1984; Arnold et al. 2001, 2008; Jones et al. 2003, 2014; Revell 2007; Melo and Marroig 2015), potentially making **G** as much as a product of adaptive evolution as a constraint upon it (Merilä and Björklund 2004).

Because stability and evolutionary lability of **G** are both plausible theoretical outcomes, the relative importance of history and adaptation in shaping **G** is largely an empirical question (Arnold et al. 2008). Several studies have shown a relationship between the shape of **G** and divergence, suggesting the importance of genetic constraints in channeling evolutionary outcomes (Bégin and Roff 2003; 2004; Blows and Hoffmann 2005; McGuigan et al. 2005; McGuigan 2006; Hansen and Houle 2008; Walsh and Blows 2009; Chenoweth et al. 2010; Bolstad et al. 2014; Houle et al. 2017; McGlothlin et al. 2018a; Walter et al. 2018). The early stages of adaptive radiation are expected to be aligned with the “genetic line of least resistance” describing the direction of greatest genetic variation within a population (Schluter 1996).

Although natural selection should be able to push phenotypes away from this line given enough time, recent results suggest that evolutionary change may be predicted by axes of genetic variation for tens of millions of years (Houle et al. 2017; McGlothlin et al. 2018a). Conversely, it is well established that both directional and nonlinear selection may alter aspects of **G**. Using an individual-based model, Melo and Marroig (2015) showed that directional selection may favor a reorientation of **G** in ways that facilitate evolutionary response. In general, traits selected in the same direction are expected to evolve stronger correlations, while traits selected in opposite directions should evolve weaker correlations. In support of these predictions, studies in natural populations of chipmunks (Assis et al. 2016) and laboratory mice (Penna et al. 2017) have demonstrated the evolution of stronger phenotypic covariance in response to concordant directional selection. Correlational selection, a form of nonlinear selection that occurs when certain combinations of traits are favored over others, can also directly alter the strength of genetic correlations (Phillips and Arnold 1989; Jones et al. 2003; Revell 2007). Patterns of genetic correlation are often congruent with axes of correlational selection in the wild (Brodie 1989; 1992; McGlothlin et al. 2005; Roff and Fairbairn 2012), and genetic correlations have been shown to evolve in response to artificial correlational selection (Delph et al. 2011; Steven et al. 2020).

Although comparative studies of **G** have become more common in recent years (Steppan et al. 2002; Bégin and Roff 2003; Hine et al. 2009; Eroukhmanoff and Svensson 2011; Walter et al. 2018), none have been able to disentangle the effects of shared ancestry from similar selection pressures in determining the evolution of **G**. Convergent evolution of **G** or its elements in similar environments would provide strong evidence that changes in **G** represent adaptation of genetic architecture (Losos 2011). The adaptive radiation of West Indian *Anolis* lizards provides an ideal testing ground for hypotheses about the evolution of **G** because the effects of phylogenetic history and ecological selection are largely decoupled (Losos 1994; 2009; 2011). In the Greater Antilles, anoles have diversified into 120 species, 95 of which can be classified as one of six habitat specialists, or ecomorphs, each of which has evolved multiple times throughout the *Anolis* radiation (Williams 1972; Losos et al. 1998; Beuttell and Losos 1999; Losos 2009). Species with dissimilar morphology on the same island tend to be more closely related than are those with similar morphology on different islands, indicating that the characteristic morphology of ecomorphs is due to convergent evolution (Losos et al. 1998; Harmon et al. 2005; Mahler et al. 2013). This repeated adaptive radiation leads to explicit predictions for the evolution of **G**. If **G** responds predictably to similar selection pressures, **G** should show signatures of convergence among the independent origins of the same ecomorph. Conversely, if **G** evolves relatively slowly and does not respond predictably to similar selection pressures, **G** or its elements should be more similar within lineages than within ecomorph classes.

Previous work in *Anolis* using phenotypic variance-covariance matrices (**P**) as proxies for **G** (Cheverud 1988) suggests that selection may indeed lead to convergence in (co)variance structure. A study comparing **P** in eight *Anolis* species showed significant variation in covariance structure across the radiation and demonstrated convergent changes in **P** in three distantly related species from the same ecomorph class (Kolbe et al. 2011). In a separate study, **P** showed significant alignment with the matrix of nonlinear selection (**γ**) in *A*. *cristatellus*, suggesting that contemporary stabilizing and correlational selection may act to shape the pattern of phenotypic (co)variance within species (Revell et al. 2010). These results suggest that selection plays a role in shaping genetic architecture in anoles, but patterns of phenotypic covariance do not necessarily mirror patterns of genetic covariance (Hadfield et al. 2007). Thus, comparative studies that directly estimate **G** are necessary to test whether its structure is more influenced by phylogenetic history or convergent evolution.

In this study, we compare **G** matrices in seven *Anolis* species reared in a controlled laboratory environment. We chose species from lineages originating on three different islands, Puerto Rico, Jamaica, and Cuba, and included three ecomorphs, trunk-crown (three species), trunk-ground (three species), and grass-bush (one species), which are distinguished by their habitat use, coloration, and skeletal morphology (Williams 1972; Beuttell and Losos 1999; Harmon et al. 2005; Losos 2009). Trunk-crown lizards are typically found high in trees and are usually green with relatively short legs for climbing and clinging to narrow perches. Trunk-ground lizards tend to be found on low perches or on the ground and are typically brown with long hindlimbs that aid in running quickly and jumping far (Losos and Sinervo 1989; Losos 1990; Irschick and Losos 1998; Beuttell and Losos 1999). The third ecomorph, grass-bush, has a slender body that matches its narrow perches and long hindlimbs that allow it to both run and jump well (Losos 1990; Beuttell and Losos 1999). The three trunk-crown species are distantly related to one another, as are the trunk-ground species. Both ecomorphs may have evolved three separate times, although it cannot be ruled out that one of these ecomorphs represents the ancestral state for the anole radiation (Losos 2009). Because of the importance of skeletal morphology, and limb length in particular, to the evolution of these ecomorphs, our estimates of **G** focus on skeletal traits.

Our previous results have shown that **G** varies substantially across these *Anolis* species, while retaining conserved axes of genetic variation (McGlothlin et al. 2018a). Specifically, **G** matrices varied most in size (overall genetic variance), and the major axis of genetic variance remained similar in orientation across all species. This major axis of genetic variance was similar in orientation to the major axis of morphological divergence, suggesting that divergence has occurred along a genetic line of least resistance even though **G** has not remained constant. The largest evolutionary changes in **G** were also aligned with the major axes of both genetic variance and morphological divergence. This triple alignment may have been caused by deep genetic constraints, natural selection, genetic drift, or some combination of the three (McGlothlin et al. 2018a).

Here, we explicitly consider the role of selection in shaping **G**-matrix evolution across the *Anolis* radiation by testing whether aspects of **G** show patterns of convergence that mirror the repeated evolution of ecomorphs. To do so, we use two types of comparisons. First, we test for convergence of the overall structure of **G** by asking whether random skewers correlations, which are estimates of pairwise similarity in the predicted multivariate response to selection, are better predicted by shared evolutionary history or shared ecology. Second, we conduct similar tests for individual elements of **G** (i.e., variances and covariances of individual traits) to test for signatures of convergence on a finer scale. Although many processes, including both selection and drift, could lead to similarities in **G** among more closely related species, convergence in the structure of **G** among distantly related species of the same ecomorph would provide strong evidence that **G** may be predictably shaped by common selection pressures.

## Methods

### Estimation of **G**

Detailed methods for estimation of the **G** matrices used here are reported elsewhere (McGlothlin et al. 2018a). Briefly, adults from seven *Anolis* species, representing independent origins of trunk- crown (*A. evermanni*, Puerto Rico; *A. grahami*, Jamaica; *A. smaragdinus*, a Bahamian species descended from *A. porcatus* on Cuba), trunk-ground (*A. cristatellus*, Puerto Rico; *A. lineatopus*, Jamaica; *A. sagrei*, Cuba), and grass-bush ecomorphs (*A. pulchellus,* Puerto Rico), were collected from the wild (fig. 1; table 1). Due to travel restrictions, species from Cuban lineages were collected from South Bimini, The Bahamas, where they occur naturally. These seven species shared a common ancestor approximately 41.5–43.5 million years ago, and the most recent phylogenetic split (between *A. cristatellus* and *A. pulchellus*) dates is estimated at 19.8–22.5 million years ago (fig. 1, Zheng and Wiens 2016; Poe et al. 2017).

**Figure 1:**
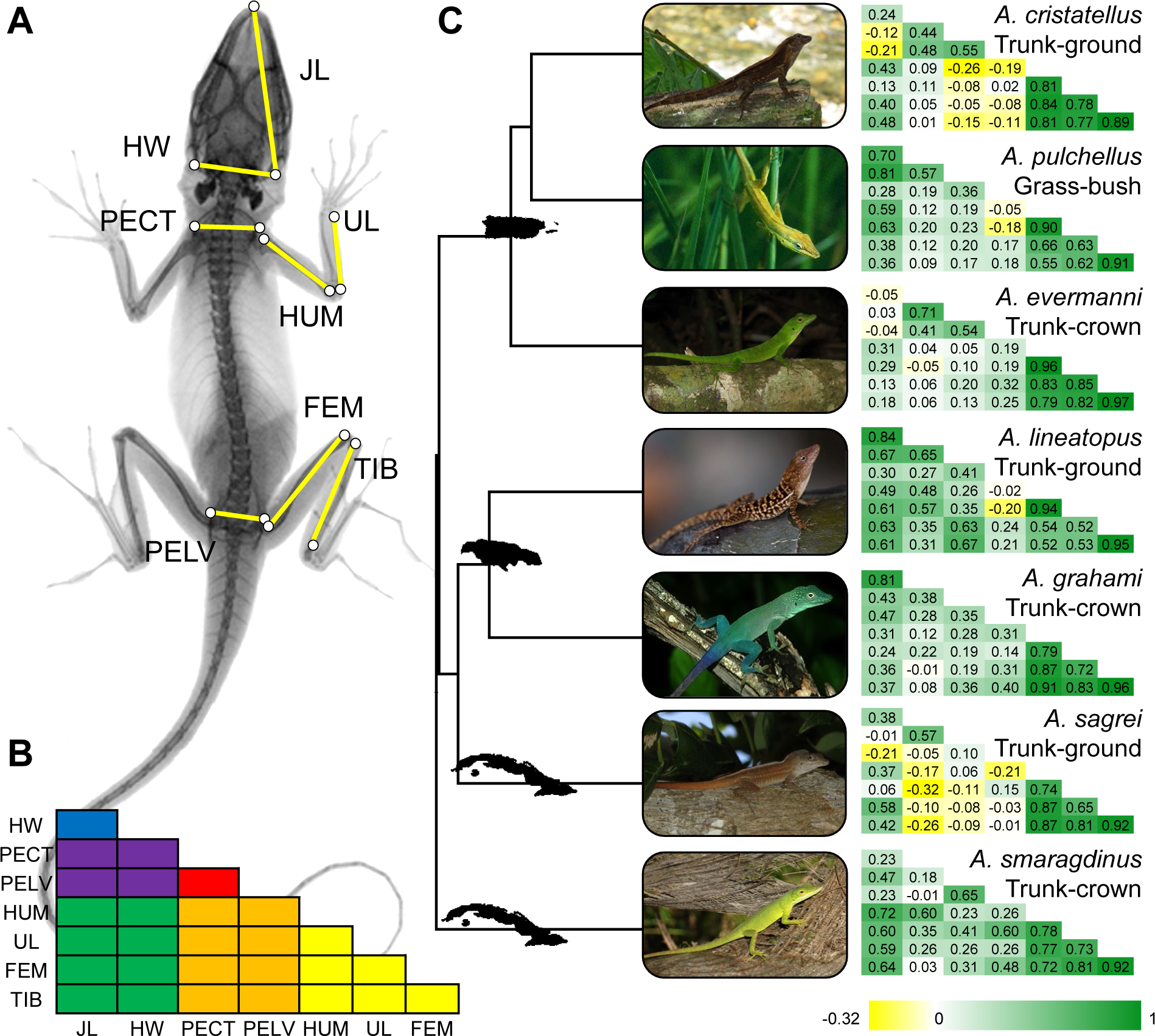
(*A*) Traits measured in this study: JL = jaw length, HW = head width, PECT = pectoral width, PELV = pelvis width, HUM = humerus, UL = ulna, FEM = femur, TIB = tibia. (*B*) Schematic of a genetic correlation matrix showing the location of each trait. Elements are color- coded based on phenotypic groups (head, body, and limbs; see Methods), with within-group correlations in primary colors (blue = head, red = body, yellow = limbs) and between-group correlations in secondary colors (violet = head-body, green = head-limb, orange = body-limb). (*C*) Genetic correlation matrices for each species. Positive correlations are shown in green and negative correlations are shown in yellow, with brighter colors signifying stronger correlations. Photographs by J.B.L. (*A*. *pulchellus*, *A*. *lineatopus*, and *A*. *graham*i), and E.D.B. III (all other species).

**Table 1:** Study design (reproduced from McGlothlin et al. 2018a)

| Species name | Ecomorph | Island | Coordinates | Sires | Dams | Juveniles |
| --- | --- | --- | --- | --- | --- | --- |
| *A. cristatellus* | Trunk-ground | Puerto Rico | 18.05°N,<br>65.83°W | 67 | 109 | 643 |
| *A. pulchellus* | Grass-bush | Puerto Rico | 18.26°N,<br>65.71°W | 35 | 62 | 430 |
| *A. evermanni* | Trunk-crown | Puerto Rico | 18.27°N,<br>65.72°W | 68 | 105 | 469 |
| *A. lineatopus* | Trunk-ground | Jamaica | 18.32°N,<br>76.81°W | 30 | 42 | 259 |
| *A. grahami* | Trunk-crown | Jamaica | 18.32°N,<br>76.81°W | 32 | 35 | 144 |
| *A. sagrei* | Trunk-ground | South Bimini,<br>The Bahamas<br>(Cuban lineage) | 25.70°N,<br>79.28°W | 55 | 99 | 791 |
| *A. smaragdinus* | Trunk-crown | South Bimini,<br>The Bahamas<br>(Cuban lineage) | 25.70°N,<br>79.28°W | 43 | 60 | 168 |

Adults were housed in individual cages in the laboratory except when paired for breeding and held at controlled photoperiod (12L:12D for Puerto Rican and Jamaican adults and 13L:11D for Bahamian adults), temperature (28°C during the day and 25°C at night), and relative humidity (65%). Lizards were provided with a perch, a mesh hammock for basking near an adjacent UVB bulb, and a carpet substrate. Adults were mated in a paternal half-sib breeding design (average of 47 sires and 69 dams per species) to produce offspring (table 1, see McGlothlin et al. 2018a for more sampling details). Laying females were provided with potted plants, which were checked weekly for eggs, which were placed in individual cups with a 1:1 mixture of water and vermiculite and held in an incubator at 28°C and 80% humidity until hatching.

Juveniles were reared in individual cages until 6 months of age and were provided with crickets and water daily. At 0, 1, 3, and 6 months of age, we X-rayed juveniles after chilling them for 10 min at 5°C in small plastic bags. The bags were then secured with masking tape to a film cartridge (Kodak Biomax XAR) for imaging in a Faxitron 43805N radiography system.

Developed radiographs were digitized using a flatbed scanner. Using ImageJ (NIH), we measured snout-vent length (SVL), two head-shape traits, jaw length (JL) and head width (HW), two body-shape traits, pectoral width (PECT) and pelvis width (PELV), and four limb-length traits, humerus (HUM), ulna (UL), femur (FEM), and tibia (TIB). SVL was measured using ImageJ’s segmented line tool as the distance from the tip of the snout to the sacrocaudal junction (McGlothlin et al. 2018a), and other traits were measured using the straight line tool as shown in fig. 1A. In all, 9,369 individual X-ray images were measured from 2,904 lab-reared juveniles from 512 maternal families (table 1, McGlothlin et al. 2018a). We used multivariate repeated measures animal models in ASReml 3.0 (Gilmour et al. 2009) to estimate **G** matrices for natural- log transformed traits, with size (natural-log SVL) as a covariate to correct for age and growth.

These models included two random animal effects, one linked to the pedigree to estimate additive genetic (co)variance and a second unlinked effect to estimate effects of permanent environment. Most species could not be reliably sexed as juveniles; therefore, we did not correct for sex in our models. In one species that has been studied extensively in the laboratory, *A. sagrei*, sexual size dimorphism is not noticeable at hatching and only becomes elaborated after 6 months of age with the maturation of testes in males (Cox et al. 2017). Genetic correlations are shown (along with heritabilities) in table A1 and visualized in fig. 1; full **G** matrices, reprinted from McGlothlin et al. (2018a), are also shown in table A1. Permanent environment (co)variances, which were generally at least an order of magnitude smaller than genetic (co)variances, and residual (co)variances are not presented here but were used in the calculation of total phenotypic variance for calculating heritabilities. As reported previously, in all but two species, all eight traits we measured were significantly heritable (mean *h*^2^ across species ± s.d.: JL, .40 ± .150; HW, .22 ± .084; PECT, .21 ± .073; PELV, .22 ± .046; HUM, .16 ± .047; UL, .15 ± .042; FEM, .45 ± .143, TIB, .54 ± .091, table A1; McGlothlin et al. 2018). In general, genetic correlations were strong and positive for pairs of limb traits and both weaker and more variable across species for other trait combinations (fig. 1, table A1; McGlothlin et al. 2018).

### Statistical Analyses

All statistical analyses using **G** matrices were performed in R 4.1.1 (R Core Team 2021). Each **G** matrix was associated with estimation error, which is quantified by the sampling (co)variance matrix of (co)variance components, calculated by ASReml as the inverse of the average information matrix (Gilmour et al. 2009). To incorporate this estimation error into our analyses, we used the restricted maximum-likelihood multivariate normal (REML-MVN) method developed by Houle and Meyer (2015). The REML-MVN method uses point estimates and their associated error distribution to define a multivariate normal distribution, from which a large number of samples are drawn. These samples may be then be used to perform a calculation, providing a distribution of output values that incorporates estimation error of the original estimates. We used point estimates of **G** and the associated sampling (co)variance matrix to define a multivariate normal distribution for REML-MVN sampling. From this distribution, we generated 10,000 samples of **G** for each species using the function rmvn from the R package mgcv 1.8-36 (Wood 2012). This set of 10,000 samples was then used to perform downstream calculations. For each analysis, we report parameter estimates from both the point estimates of **G** and from the median of the distribution resulting from performing the analysis using the set of 10,000 REML-MVN samples. The distribution of REML-MVN was also used to calculate 95% confidence intervals, which we report as the 2.5% and 97.5% quantiles. A parameter estimate was considered to be statistically supported when the expected null value (often zero) was outside of this 95% confidence interval.

To quantify pairwise similarity in the overall structure of **G**, we used random skewers analysis, which compares the response to selection predicted by a pair of **G** matrices (Marroig and Cheverud 2001; Cheverud and Marroig 2007; Revell 2007; Aguirre et al. 2014). Random skewers analysis has the advantage of providing an evolutionarily relevant comparison of **G** using a single metric. We used the function RandomSkewers in the R package evolQG 0.2-9 (Melo et al. 2015) to apply 10,000 random selection gradients (**β**) to all seven **G** matrices. Each element of these skewers was drawn from a normal distribution, after which each skewer was normalized to unit length and used to calculate the predicted multivariate selection response (Δ*z̄*) from the multivariate breeder’s equation, Δ*z̄* = Gβ (Lande 1979). The correlation in response to selection for two species, or the random skewers correlation (*r*_RS_), was calculated as the pairwise vector correlation of the resultant 10,000 estimates of Δ*z̄*. To incorporate estimation error, *r*_RS_ was recalculated for each of 10,000 sets of seven **G** matrices each generated from REML-MVN resampling. We report the median and 95% confidence intervals of this distribution in addition to our point estimate. We found that incorporating error from pairs of **G** matrices led to negatively skewed REML-MVN distributions of *r*_RS_. As a result, medians of the REML-MVN distributions tended to be lower than *r*_RS_ estimates calculated using point estimates of **G**.

To test for phylogenetic signal in the overall structure of **G**, we used a multivariate form of Blomberg’s *K* (Blomberg et al. 2003), *K*_mult_, developed by Adams (2014) and implemented in the R package geomorph 4.0.0 (Adams and Otárola-Castillo 2013). In general, Blomberg’s *K* estimates phylogenetic signal by measuring deviations from Brownian motion model of evolution. The expected value of *K* under Brownian motion is 1, with *K* < 1 indicating more similarity in distantly related species and *K* > 1 indicating greater than expected similarity in closely related species. When the phylogeny provides no information about trait distribution, the expected value of *K* is 0. Therefore, we tested for significant differences from both *K*_mult_ = 0 (no phylogenetic signal) and *K*_mult_ = 1 (phylogenetic signal consistent with Brownian motion). We calculated *K*_mult_ for random skewers correlations following Machado et al. (2018). We first converted each value of *r*_RS_ to a distance using the formula 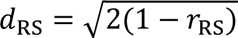. We then calculated the principal coordinates of the distance matrix of *d*_RS_ values. The resulting vectors, along with a tree pruned from a dated squamate phylogeny (Zheng and Wiens 2016) were used to calculate *K*_mult_ with the function physignal in geomorph. Dates from a tree of all *Anolis* species (Poe et al. 2017), which had identical topology for our species, were similar. We calculated *K*_mult_ separately using our point estimates of **G** and our REML-MVN samples, allowing us to present a value of *K*_mult_ with 95% confidence intervals.

To test for similarity of **G** between species of the same ecomorph, we used a Mantel test with phylogenetically informed permutations (Lapointe and Garland 2001; Harmon and Glor 2010) implemented using the function PhyloMantel in evolQG to compare the distance matrix of *d*_RS_ values to an ecomorph distance matrix, where 0 was used to represent species of the same ecomorph and 1 was used to represent species of different ecomorph. This test was conducted using only trunk-ground and trunk-crown species, a phylogeny pruned to six species, and the phylogeny parameter *k* set to the default value of 1. A positive Mantel correlation (*r*_M_ > 1) would indicate convergence, i.e., that species from the same ecomorph tend to have more similar **G** matrices. In addition to the phylogenetically informed Mantel test, which we performed only using our point estimates of **G**, we report results from standard distance matrix correlations of our REML-MVN replicates, which allows us to report *r*_M_ with 95% confidence intervals.

Evolutionary patterns in **G** may involve changes that are subtler than can be detected in analyses of its overall structure. Therefore, we also tested for the effects of shared evolutionary history and shared ecology on the individual elements of **G**. We compared genetic variances (diagonal elements of **G**) and genetic correlations (off-diagonal elements of **G** standardized by the square root of the product of the variances) across **G** matrices, testing for both phylogenetic signal and differences between ecomorphs (trunk-crown vs. trunk-ground). We present comparisons of genetic correlations rather than genetic covariances so that tests for associations between traits would be independent of differences in variance across species. However, we note that analyses using covariances gave nearly identical results (not shown). Because REML-MVN occasionally produced non-positive definite matrices, we produced 10,000 samples from separate univariate normal distributions defined by genetic correlations and their approximate standard errors (table A1) to incorporate estimation error into analyses that used genetic correlations.

To test for the effects of shared evolutionary history, we estimated Blomberg’s *K* for each genetic variance and genetic correlation using the R package phytools 0.7-80 (Revell 2012). We tested for effects of shared ecology on genetic variance and genetic correlations of trunk-ground and trunk-species using phylogenetic generalized least squares (Martins and Hansen 1997) with ecomorph as a predictor (coding trunk-crown as 0 and trunk-ground as 1; *A*. *pulchellus* was excluded), our dated tree, and an assumption of Brownian motion evolution (implemented in the R package APE, Paradis et al. 2004). Alternative models that assumed an Ornstein-Uhlenbeck evolutionary model provided similar results (not shown). For each test, we report the evolutionary correlation (*r*_e_) between the element of **G** and ecomorph to remove effects of scale.

For a given element of **G**, *r*_e_ > 0 indicates a larger value for trunk-ground species, and *r*_e_ < 0 indicates a larger value for trunk-crown species. Both of these analyses were conducted using both our point estimates and our resamples to incorporate estimation error. Using the 95% confidence intervals from the resampled distributions, we note deviations from both *K* = 0 and *K* = 1 for phylogenetic signal and *r*_e_ = 0 for the evolutionary correlation.

For visualization, we grouped our estimates of phylogenetic signal and evolutionary correlation into six categories. Three categories consisted of elements of **G** within phenotypic groups, head (JW, HL), body (PECT, PELV), and limbs (HUM, UL, FEM, and TIB), and three contained elements of **G** between phenotypic groups, head-limb, head-body, and body-limb.

Because our sample sizes differed across **G** matrices, one possible concern is that our results may be an artifact of variation in sample size. To test for this possibility, we examined a reduced data set where each **G** matrix was estimated using a small subsample of the data (21–25 sires, 31–35 dams, and 104–144 juveniles). We found that these results were consistent with those from the full sample (results not shown), suggesting that the results reported below were not statistical artifacts.

## Results

### Overall Structure of **G**

Predicted responses to selection were highly correlated between all pairs of species (median *r*_RS_ =, range .75–.92, REML-MVN median *r*_RS_ = .75, range .62–.86; table A2, fig. 2), suggesting that all species have **G** matrices with similar overall structure despite being separated for 20–44 million years. **G**-matrices showed phylogenetic signal consistent with a model of Brownian motion (*K*_mult_ = .94; REML-MVN estimate: *K*_mult_ [95% CI] = .96 [.90, 1.02]), indicating that more closely related species tend to have **G** matrices that predict a more similar evolutionary response. More distantly related pairs of species also displayed more variable values of *r*_RS_, with some distantly related pairs of species showing highly similar **G** matrices and others showing highly dissimilar **G** (fig. 2). Overall similarity of **G** was not predicted by ecomorph (*r*_M_ = -.29, *p* = .96; REML-MVN estimate: *r*_M_ [95% CI] = -.10 [-.31, .14]). Despite the lack of a consistent ecomorph effect, the two distantly related trunk-ground species *A*. *cristatellus* and *A*. *sagrei* did have highly similar **G** matrices (figs. 2 & 3).

**Figure 2:**
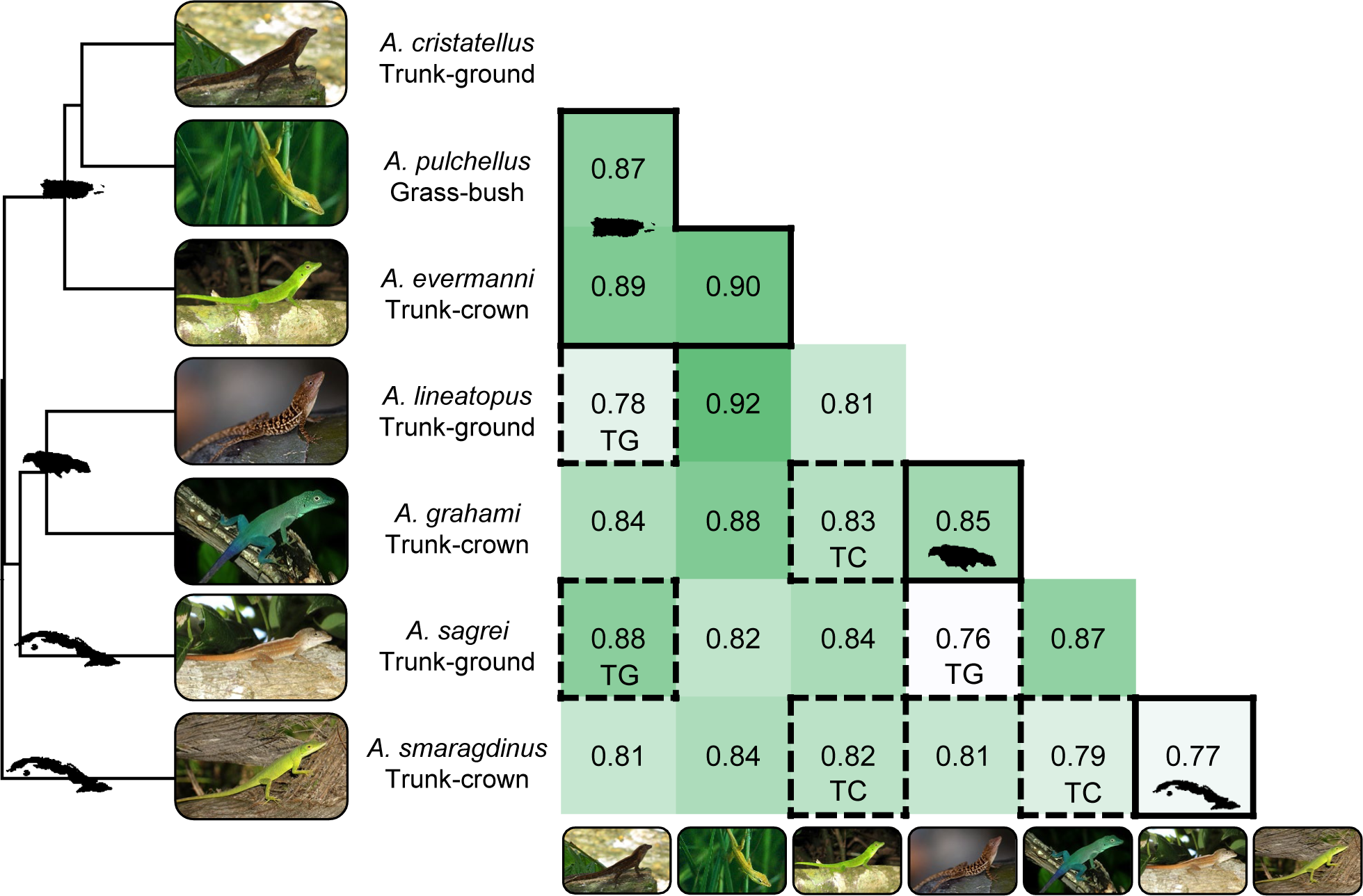
Correlations between species in the overall structure of the **G** matrix as measured via random skewers correlations (*r*_RS_). Stronger correlations are represented by darker shading. Within-island comparisons are shown with a solid border, and within-ecomorph comparisons are shown with a dashed border (TC = trunk-crown, TG = trunk-ground). Islands of origin are represented by their shapes on the phylogeny (Puerto Rico, Jamaica, and Cuba, from top to bottom) and for the three within-island comparisons. Note that despite originating on Cuba, *A. sagrei* is actually more closely related to Jamaican species than to *A. smaragdinus*. The phylogeny is pruned from a dated tree of squamates (Zheng and Wiens 2016).

**Figure 3:**
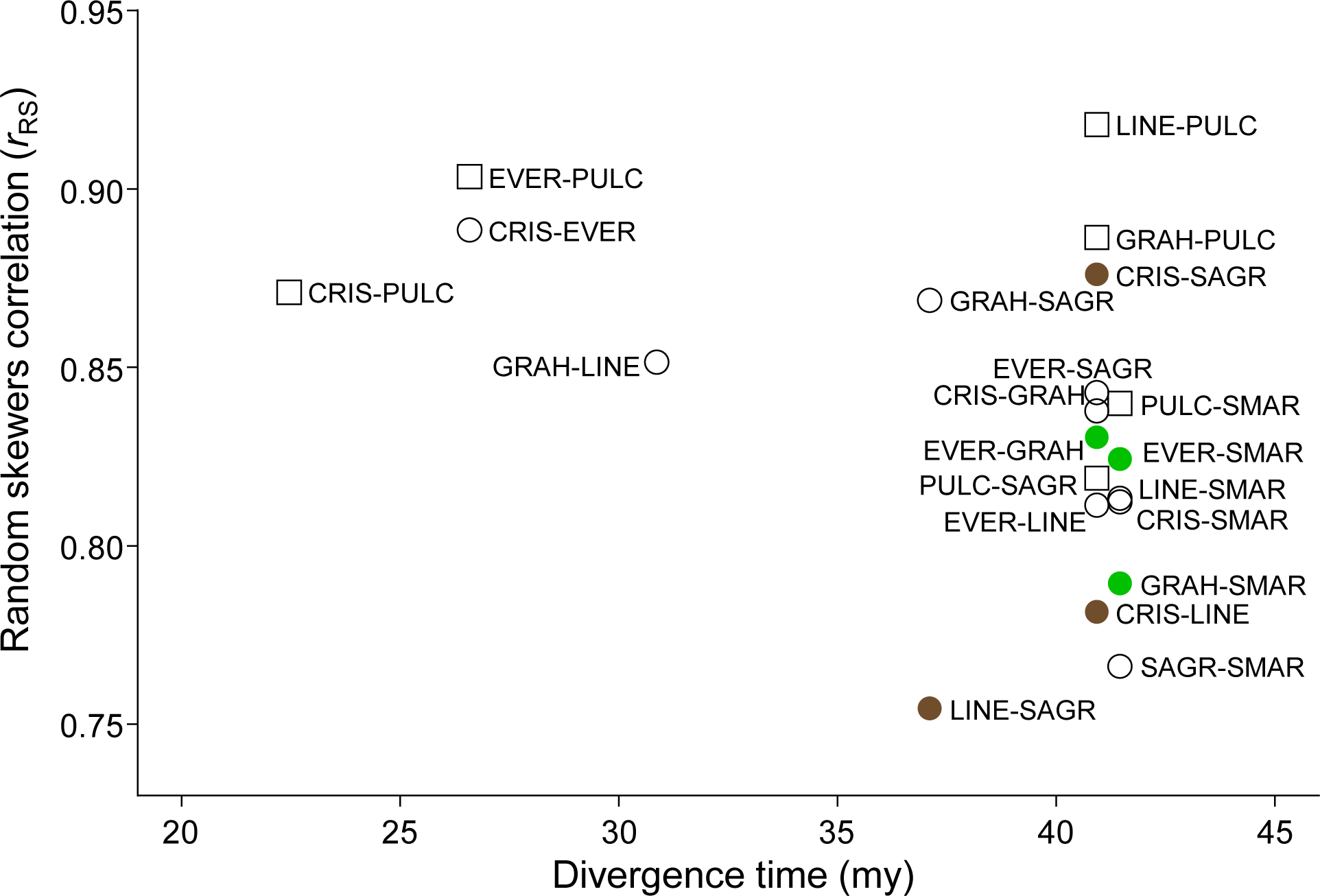
Relationship between divergence time (millions of years, my) and random skewers correlations (*r*_RS_). **G** matrices of more distantly related species are significantly less similar and more variable in *r*_RS_. Within-ecomorph comparisons are shown as colored circles (green = trunk-crown and brown = trunk-ground), with trunk-crown/trunk-ground comparisons as open circles and comparisons with the grass-bush species as open squares. Each point is labeled with four letter codes for the two species under comparison (CRIS = *A*. *cristatellus*, EVER = *A*. *evermanni*, GRAH = *A*. *grahami*, LINE = *A*. *lineatopus*, PULC = *A. pulchellus*, SAGR = *A. sagrei*, SMAR = *A. smaragdinus*).

### Individual Elements of **G**

All individual elements of **G** showed phylogenetic signal significantly higher than *K* = 0 but indistinguishable from *K* = 1, which is the null expectation of the Brownian motion model (median *K =* .91, REML-MVN estimate: *K* = .94; fig. 4A, table A3). Phylogenetic signal did not exhibit any detectable patterns across trait groups (fig. 4A).

**Figure 4.**
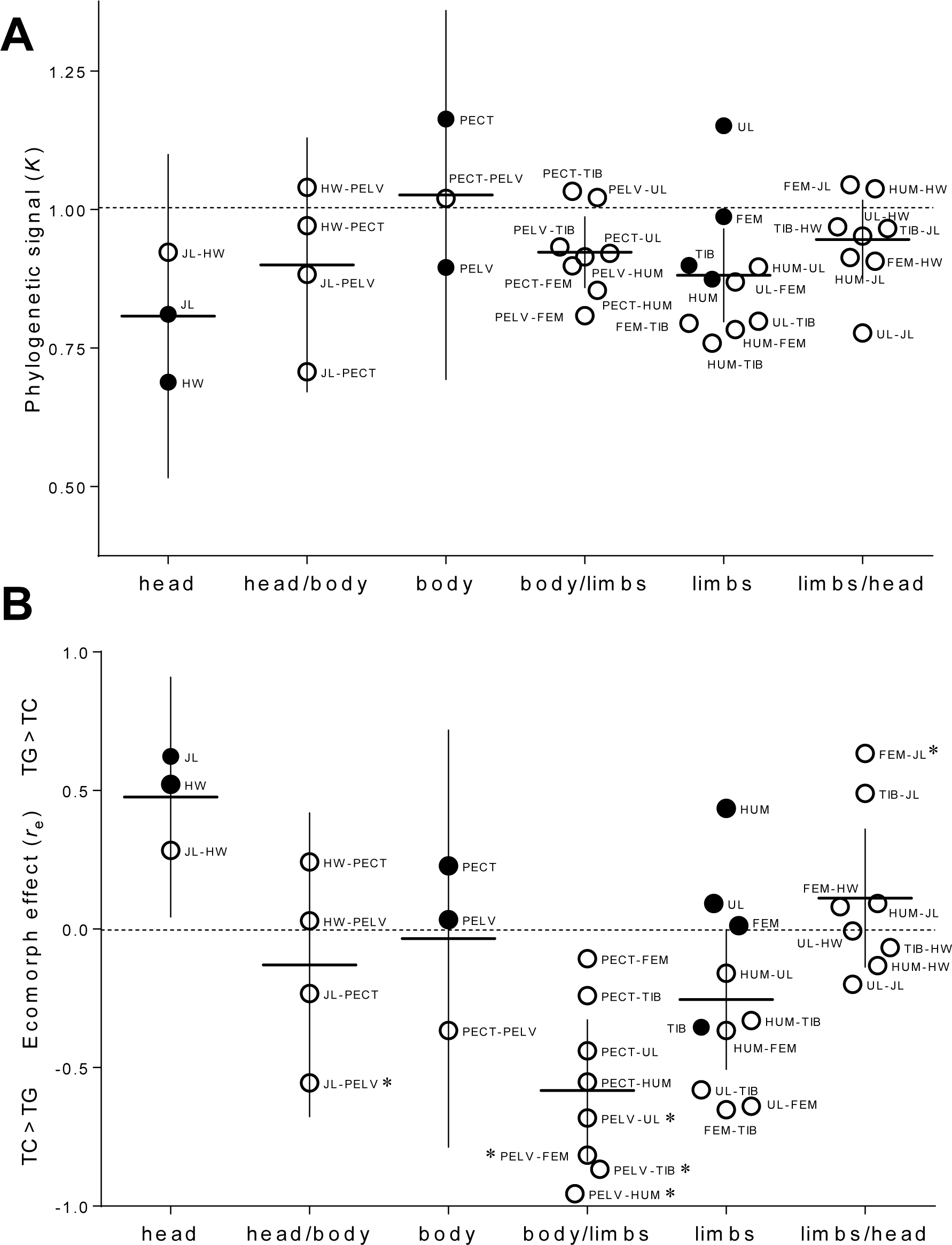
(previous page): Element-by-element comparisons of genetic variances (solid circles) and correlations (open circles). Elements are split into three within-group categories (head, body, and limbs) and three between-group categories (head/body, body/limb, and limb/head) and are labeled with abbreviations as in Fig. 1. Bars show means and 95% confidence intervals for a set of point estimates within a group and are presented for visualization purposes only. (*A*) Strength of phylogenetic signal, as estimated using Blomberg’s *K*. Estimates were clustered around *K* = 1, suggesting phylogenetic signal consistent with Brownian motion. (*B*) Ecomorph effects from phylogenetic least squares, given as the evolutionary correlation (*r*_e_). Points above the midline indicate that trunk-ground species had higher values of a given element of **G** than did trunk-crown species; the converse is true below the midline. Six genetic correlations showed a significant correlation with ecomorph (trunk-ground vs. trunk-crown; *p* < .05, denoted by *).

Six elements of **G** differed significantly between trunk-crown and trunk-ground ecomorphs. Specifically, genetic correlations between pelvis width and both jaw length and all four limb bones were significantly lower in trunk-ground species than in trunk-crown species (fig. 4B; table A4). In addition, genetic correlations between jaw length and femur length were higher in trunk-ground species than in trunk-crown species (fig. 4B; table A4). In some cases, these effects involved changes in the sign of genetic correlations from positive in trunk-crown species to negative in trunk-ground species (fig. 1). Although not a statistically significant effect, genetic correlations both among limb bones and between pectoral width and all four limb bones tended to be weaker in trunk-ground species than in trunk-crown species, indicating a trend toward weaker integration among limb and body traits in trunk-ground species.

Although we did not conduct formal tests for the lone grass-bush species, *A*. *pulchellus*, inspecting its genetic correlation matrix (fig. 1, table A1) shows that this species displays some similarities to trunk-ground lizards. In particular, this species had weak (and occasionally negative) genetic correlations between the pelvis and all limb bones, similar to all trunk-ground species. *Anolis pulchellus* also showed genetic correlations between hindlimb and forelimb bones that were noticeably weaker than the other two species in the Puerto Rican lineage (fig. 1, table A1).

## Discussion

Here we present two significant findings about the evolution of quantitative genetic architecture within the adaptive radiation of West Indian *Anolis* lizards. First, when viewed in terms of its effects on multivariate response to selection, we show that evolution of the overall structure of the *Anolis* **G** matrix exhibits a phylogenetic signal consistent with a Brownian motion model of evolution. Although the seven species we studied have been separated for 20–44 million years, all **G** matrices predicted a similar multivariate evolutionary response, and more closely related species had more similar **G** matrices. Second, despite this phylogenetic signal in the overall structure of **G**, pairwise genetic correlations between limb traits and body traits showed signatures of convergence, suggesting that they may have been adaptively shaped by similar selection pressures resulting from each ecomorph’s niche. In particular, longer-limbed trunk-ground lizards show a decoupling of limb length and pelvis width relative to shorter-limbed trunk-crown lizards, demonstrating that convergent changes in genetic architecture may accompany repeated morphological adaptation. Taken together, our results show that selection may alter **G** in predictable and evolutionarily consequential ways without leading to major changes in its overall structure.

From the perspective of overall multivariate response to selection, **G** retains similar structure across the *Anolis* radiation, with predicted responses to selection in random directions showing strong positive correlations ranging from .76 to .92. Over the span of a few generations, then, morphological divergence of species with the **G** matrices estimated here should be constrained to lie along directions defined by quantitative genetic architecture. Indeed, previous work has shown that divergence of these species remains aligned with the major axis of genetic variation, **g**_max_, even after ∼44 million years of divergence (McGlothlin et al. 2018a). The similarity of the overall structure of **G** declined with greater phylogenetic distance. This pattern suggests that the overall structure of **G** changes relatively slowly, remaining conserved over millions of years. This result suggests that multivariate stabilizing selection on the suite of traits we measured may have preserved the major features of the **G** matrix across the *Anolis* adaptive radiation (Jones et al. 2003; Melo and Marroig 2015). In addition to this phylogenetic trend, we found that more distantly related species also show greater variance in random skewers correlations. Although many distantly related species have dissimilar **G** matrices, some pairs have highly congruent **G**, a pattern that is not explained by convergent morphology. This increased variance emphasizes the unpredictability of the evolution of **G** structure over longer timescales.

When considering more subtle changes in **G**—shifts in trait-specific genetic variances and correlations—we found evidence that **G** can change repeatedly and predictably in response to similar selection pressures. The strongest convergence occurred in genetic correlations between limb bones and pelvis width, which were significantly reduced in trunk-ground species relative to trunk-crown species. In some cases, genetic correlations differed in sign between ecomorphs.

When this was the case, genetic correlations were usually negative in trunk-ground species and positive in trunk-crown species. These results indicate that the pattern of genetic integration was subtly remodeled in the transition from trunk-crown to trunk-ground ecomorphs (or vice versa), most notably in the relationship between limb length and pelvis width, for which trunk-ground species showed weaker genetic correlations when compared to trunk-crown species.

The ecomorph difference in the genetic correlations between limb length and pelvis width is likely to result from some combination of directional selection and correlational selection acting on these traits. Ecomorph differences in limb length have clear links to performance within their characteristic habitats (Losos and Sinervo 1989; Losos 1990; Irschick and Losos 1998), suggesting that they have been driven apart by divergent directional selection. The longer hindlimbs of trunk-ground lizards facilitate running faster on flatter surfaces, whereas the shorter hindlimbs of trunk-crown lizards are suited for a wider variety of perches (Losos 1990; Losos and Irschick 1996; Irschick and Losos 1998). Strong directional selection acting only on limb length could reduce its genetic correlations with other traits (Melo and Marroig 2015). It is likely that correlational selection also has played a role. The evolution of trunk-ground anoles from a hypothetical trunk-crown ancestor would require evolution of longer legs without a concomitant increase in the pelvis, which may lead to negative correlational selection to decouple the two traits. In contrast, correlational selection might favor a positive correlation between the two traits in trunk-crown anoles, perhaps because a matching limb and pelvic morphology would facilitate agility on branches.

Differences in hindlimb length between trunk-crown and trunk-ground ecomorphs are apparent at hatching and appear to emerge mostly via changes in developmental patterning early in embryonic development rather than differences in growth (Sanger et al. 2012). The developmental genetic networks underlying limb growth and development are well understood (Rabinowitz and Vokes 2012), and comparative genomic evidence indicates that genes expressed in these networks experienced enhanced positive selection during the radiation of anoles (Tollis et al. 2018). Limb-development networks share some genes in common with the network underlying development of the pelvic girdle (Sears et al. 2015). Therefore, it is likely that changes in genetic correlations between limb length and pelvis involve evolutionary changes in the expression of some of these shared genes. Future work should explore remodeling of these networks to understand the developmental genetic underpinnings of convergent morphological evolution in anoles.

Weaker genetic correlations between limbs and body traits likely facilitated the evolution of longer hindlimbs in trunk-ground anoles without correlated changes in the rest of the body.

Such genetic decoupling of limbs and body may help explain the remarkable adaptability of the trunk-ground ecomorph. Trunk-ground anoles seem to be especially capable of colonizing new habitats. Trunk-ground species have successfully colonized the Bahamas and the Virgin Islands (*A*. *sagrei* and *A*. *cristatellus*, respectively) and have become established invaders in a number of locations following introduction by humans (Kolbe et al. 2004; Eales et al. 2008; Losos 2009).

Trunk-ground anoles also have greater species richness within islands (Losos 2009), in part because they have radiated into ecologically distinct macrohabitats (Glor et al. 2003). Some of this adaptability is likely due to rapid evolution in limb length, such as has been demonstrated in experimental populations of *A. sagrei* (Losos et al. 1997, 2001; Kolbe et al. 2012). Although comparable studies have not been conducted using trunk-crown ecomorphs, their stronger genetic correlations between limb traits and body traits suggest that rapid, independent evolution of limb length would not be as likely in trunk-crown species.

The genetic correlations between pelvis and limb length in the single grass-bush anole we examined resemble those of trunk-ground anoles, suggesting either similar selection or common ancestry, or a combination of the two. Grass-bush anoles have long hindlimbs relative to their body width, suggesting that a combination of directional and correlational selection may have reduced these correlations. However, *A*. *pulchellus* is likely to have evolved from a trunk-ground ancestor (Poe et al. 2017), which suggests that both this species and the closely related *A*. *cristatellus* may have inherited weakened genetic correlations between pelvis and limbs from a common ancestor. In other respects, however, the genetic correlation structure of *A*. *pulchellus* is dissimilar to that of *A*. *cristatellus*. In contrast to *A. cristatellus*, *A. pulchellus* appears to have attained differences between the lengths of its hindlimbs and forelimbs via a reduction in the genetic correlations between the two, a feature it shares with the distantly related trunk-ground lizard *A*. *lineatopus*.

## Conclusion

The evolution of **G** in West Indian anoles illustrates how the complex interplay between selection and history influences genetic architecture. **G** reflects neither an irresistible pattern of constraint nor an easily adapted phenotype responding quickly to environmental pressures. Patterns of genetic covariation have potentially influenced the pathways that selection may follow, as evidenced by evolution along a deeply conserved genetic line of least resistance in this radiation (McGlothlin et al. 2018a). At the same time, as we show here, selection leaves an imprint, if subtle, upon **G** as species diverge. We did not observe a full-scale overhaul of **G** as ecomorphs evolved. However, small-scale differences in the elements of **G** involving links between limb length and body shape arose predictably between ecomorphs and may have had substantial evolutionary consequences. Our results emphasize that while genetic constraints may change as adaptation proceeds, these changes need not be large to facilitate phenotypic diversification.

Rather, the convergent changes we observed in the individual elements of **G**, particularly between the limbs and the body, demonstrate that consistent selection pressures can alter underlying genetic constraints in subtle ways that facilitate adaptation and influence future evolutionary potential.

## Acknowledgments

We thank Simon Pearish and Michelle Sivilich for managing the lizard colony. Dozens of undergraduates at the University of Virginia provided animal care and assistance with data collection; special thanks is due to Margo Adler, Tyler Cassidy, Brian Duggar, Maridel Fredericksen, Casey Furr, Jessie Handy, Bryan Hendrick, Uma Pendem, Jeff Wright, and Elizabeth Zipperle. Leleña Avila, Chris Feldman, Vince Formica, Tonia Hsieh, Melissa Losos, Ashli Moore, Liam Revell, and Matt Sanford assisted with field collections. We thank the editors, two anonymous reviewers, Brooke Bodensteiner, Vincent Farallo, Kerry Gendreau, Angela Hornsby, and Josef Uyeda for comments on the manuscript, and Luke Harmon, Fabio Machado, Gabriel Marroig, Emília Martins, Diogo Melo, and Liam Revell for helpful discussions. This work was supported by the National Science Foundation (grant numbers DEB 0519658 and 0650078 to E.D.B. III and DEB 0519777 and 0722475 to J.B.L.), University of Virginia, and Virginia Tech. The authors declare that there are no conflicts of interest.

## Statement of Authorship

J.B.L. and E.D.B. III conceived the study; J.W.M. and J.J.K. contributed to study design; J.W.M., J.J.K., J.B.L., and E.D.B. III performed field collections; J.W.M. and E.D.B. III oversaw the breeding experiment; J.W.M., M.E.K., and H.V.W. collected data; J.W.M. analyzed data; J.W.M. drafted the manuscript, and all authors contributed to the final version of the manuscript.

## Data and Code Availability

Raw data for estimating **G** matrices are available in a Dryad Data Repository from a previous publication (McGlothlin et al. 2018b). All code and processed data are available at https://github.com/joelmcg/AnolisG and will be archived in Dryad upon manuscript acceptance.

## Appendix

### Supplemental Tables

**Table A1:**
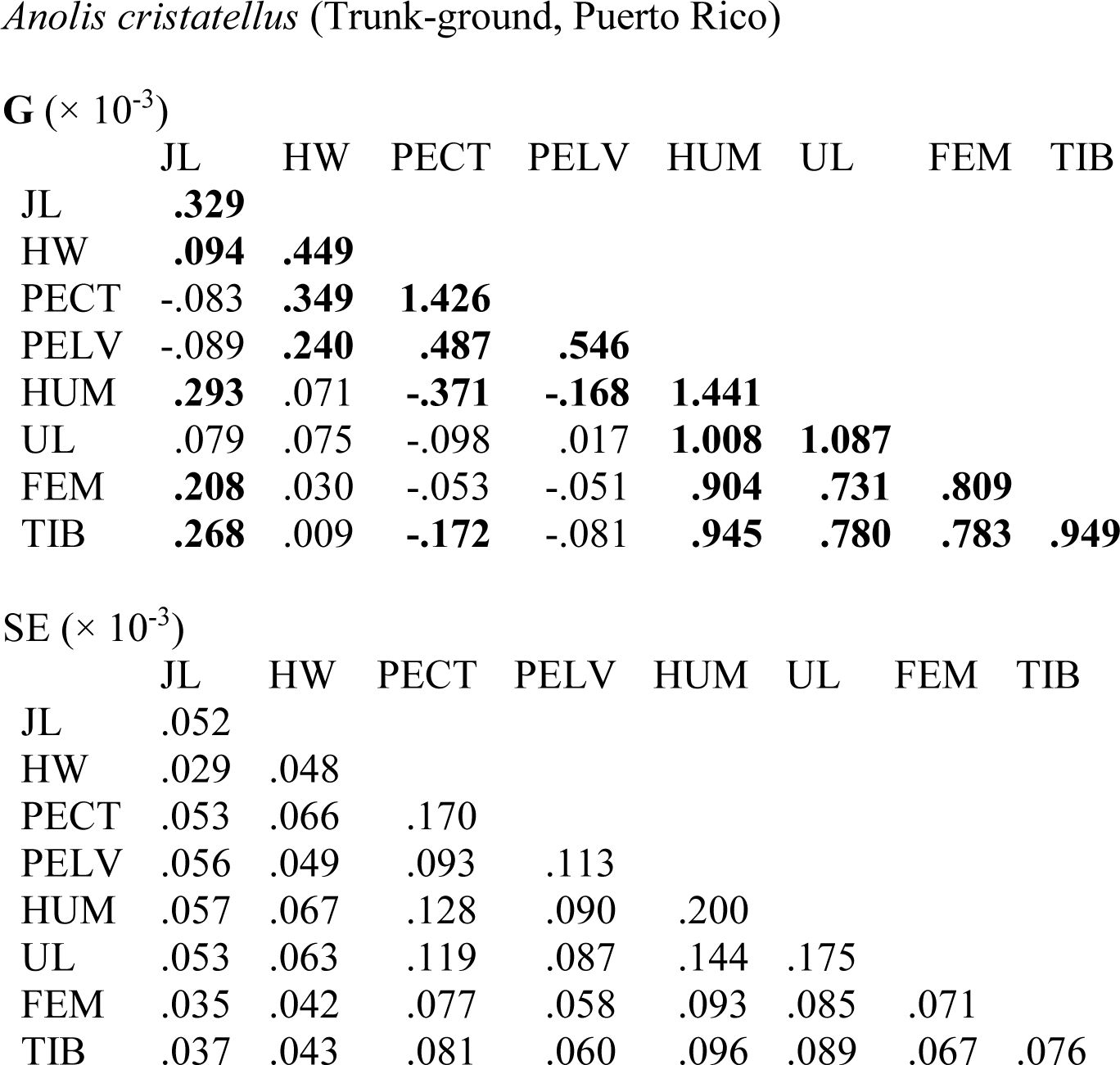
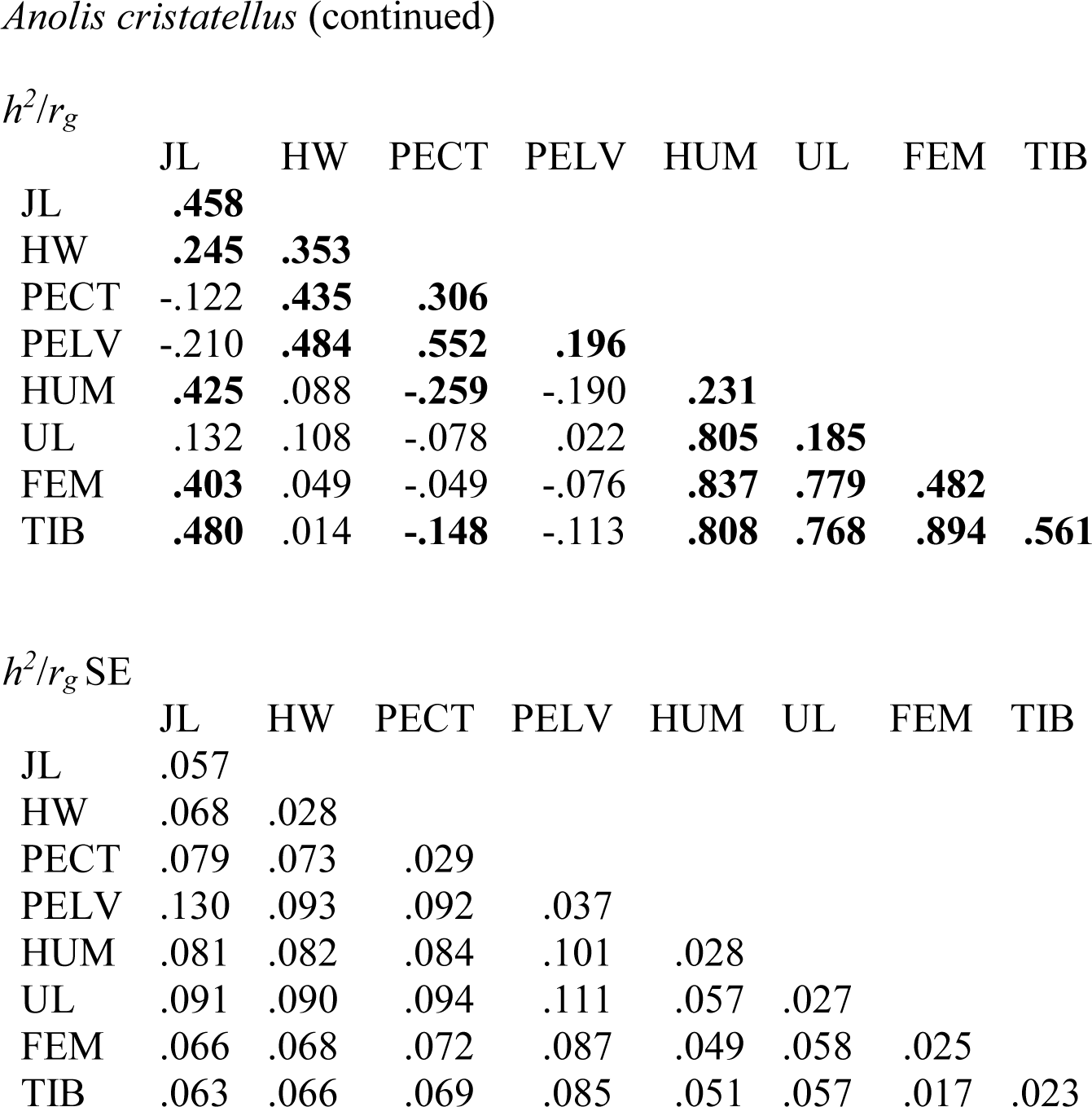
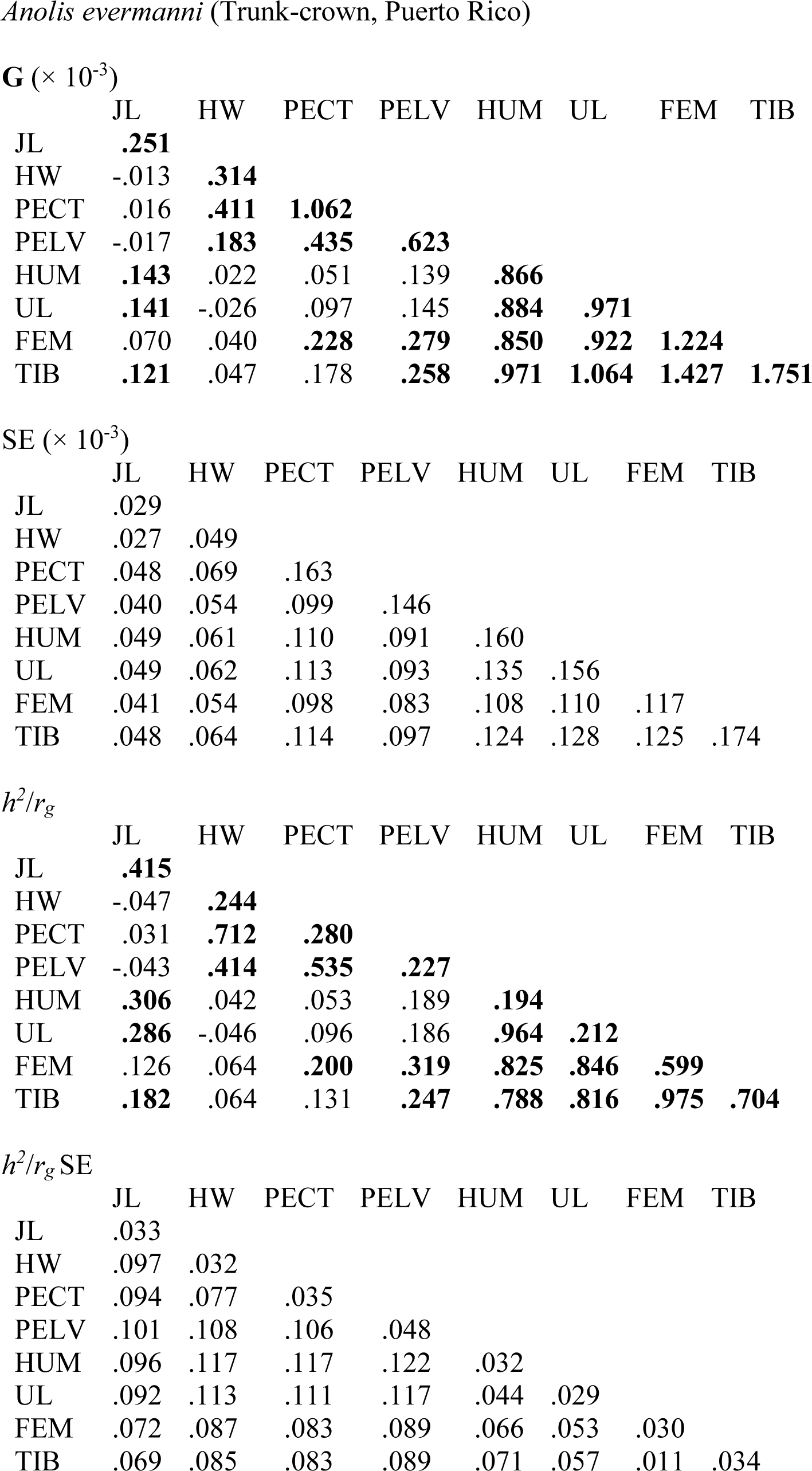
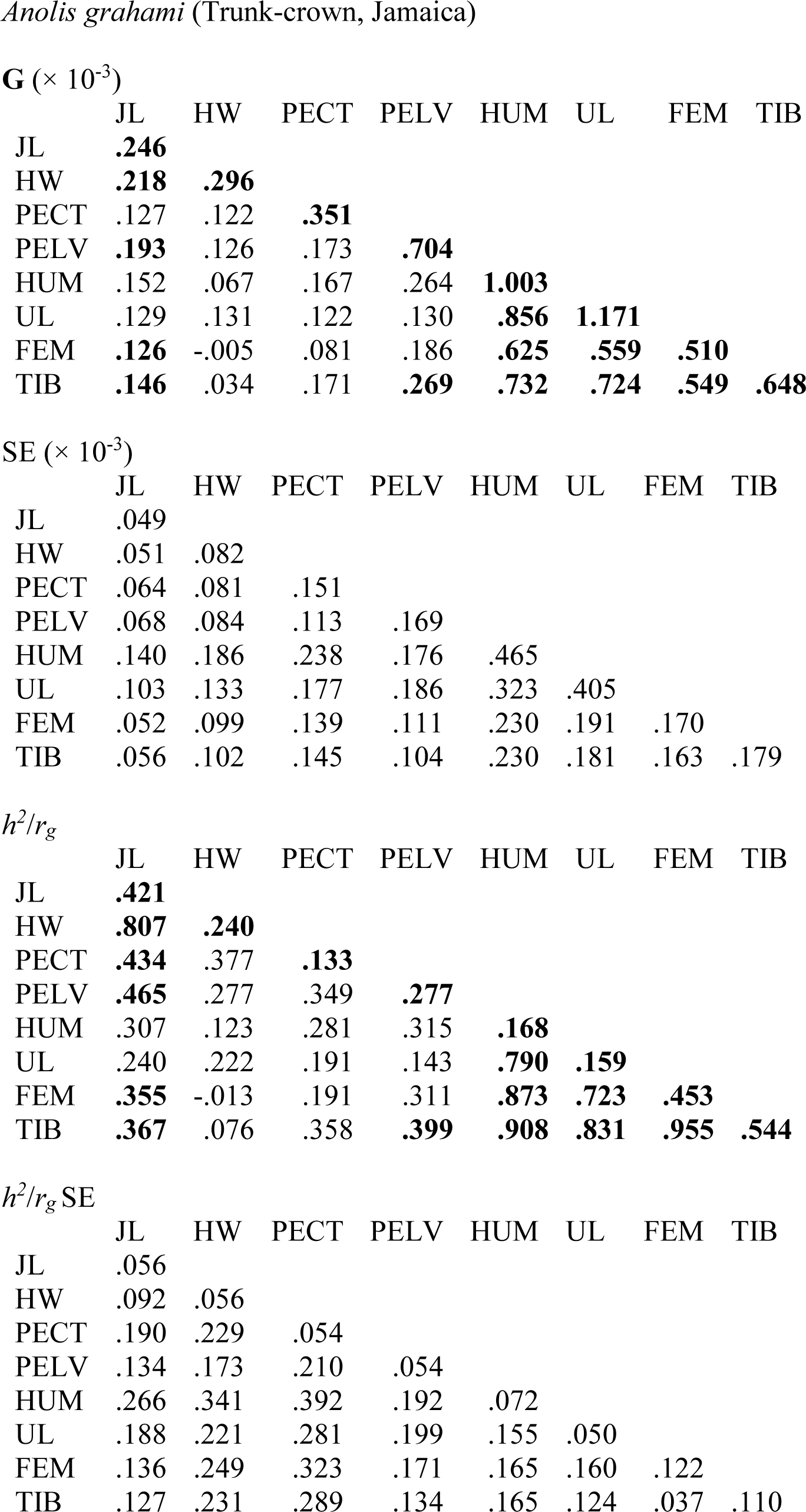
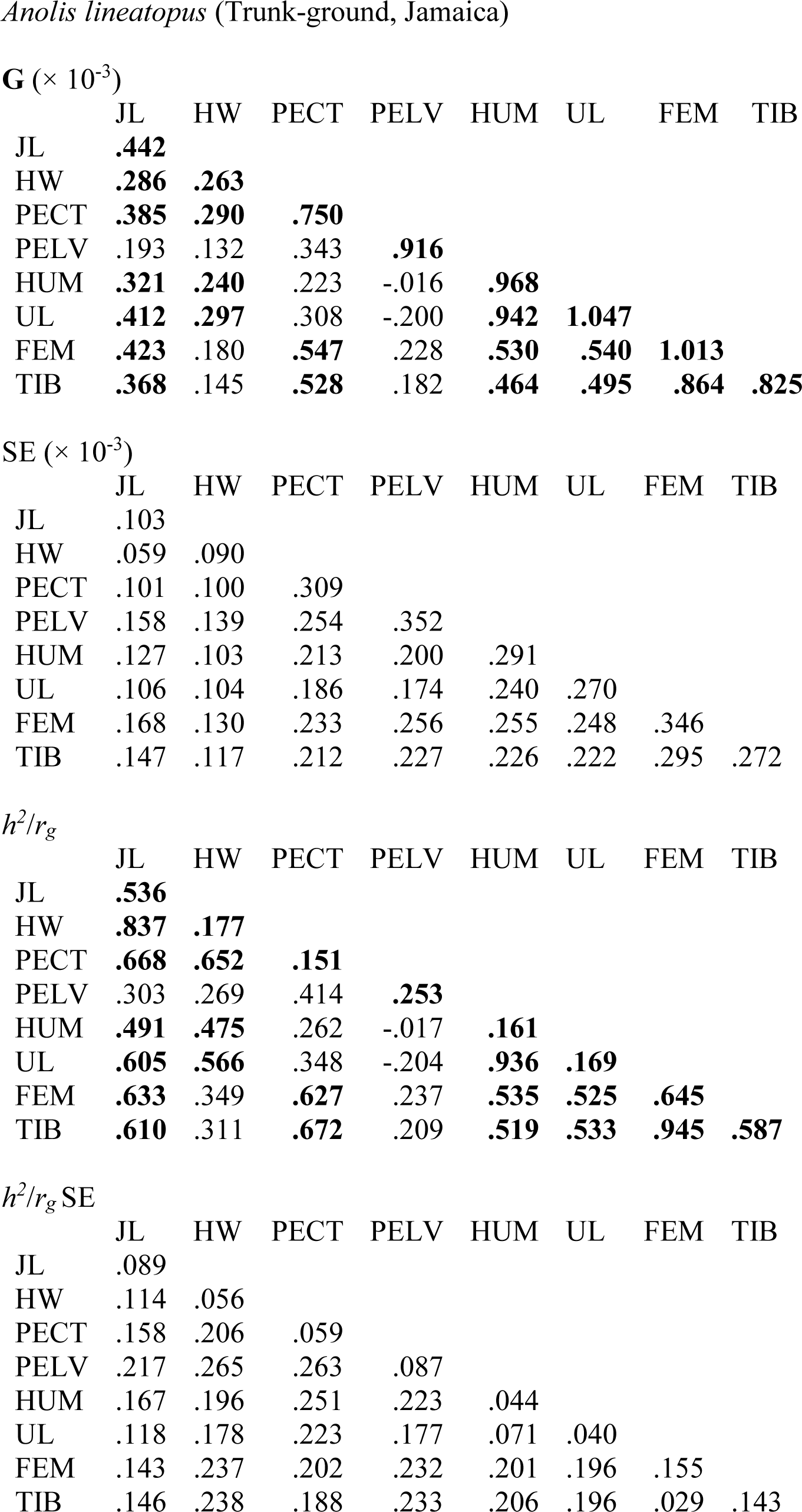
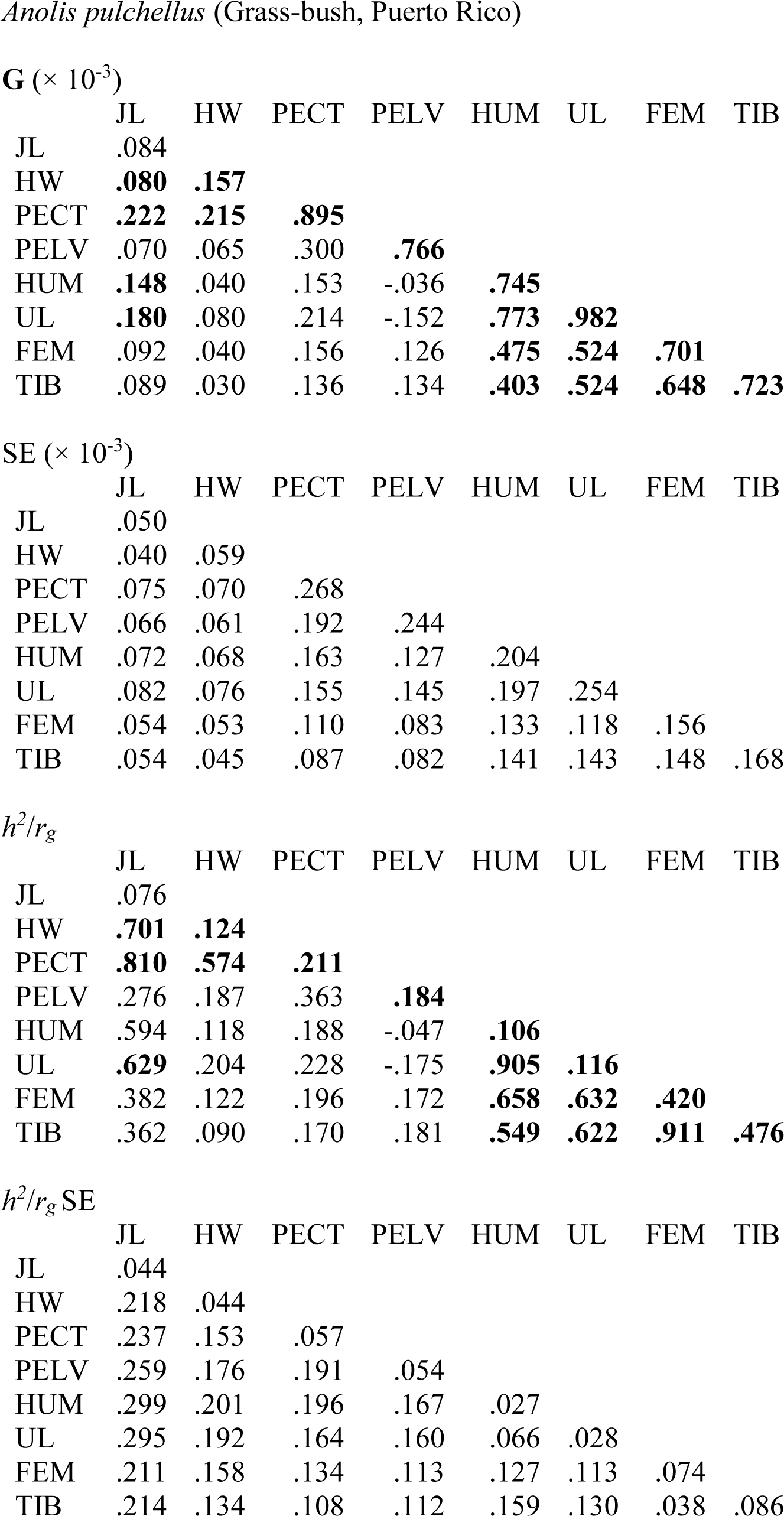
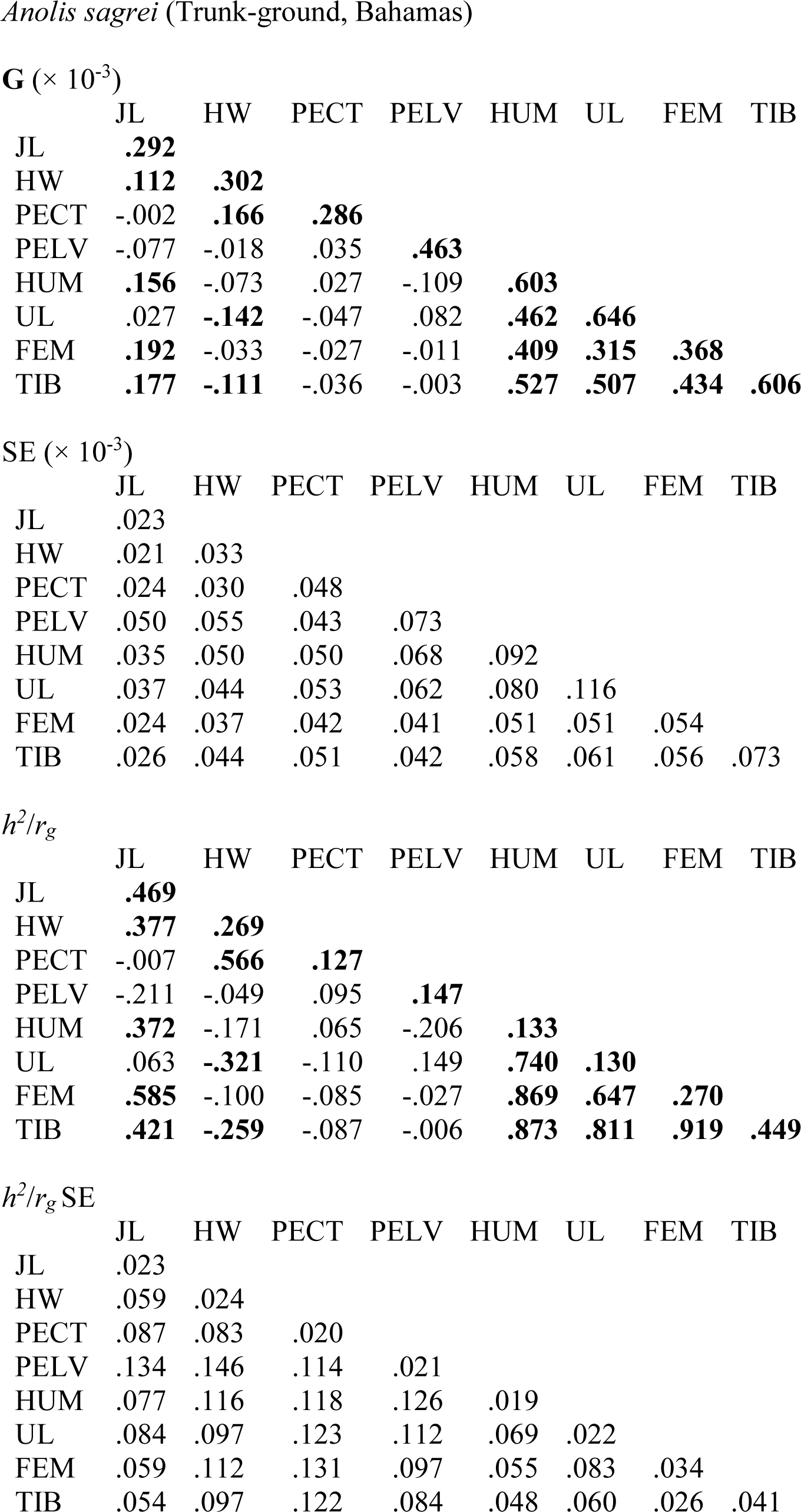
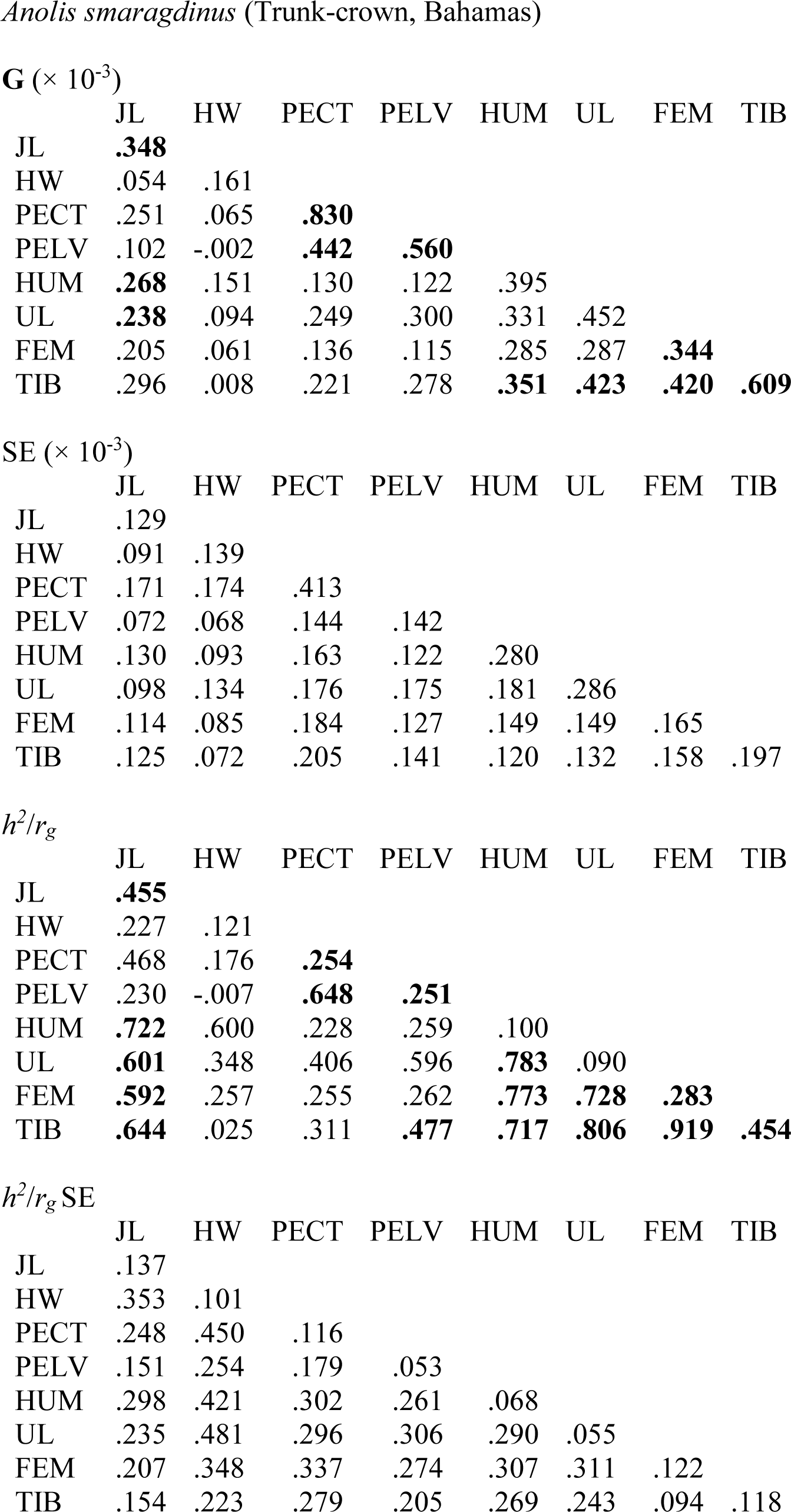
**G** matrices and matrices of heritabilities (*h*^2^, diagonal) and genetic correlations (*r_g_*, off-diagonal) for seven *Anolis* species. Approximate standard errors calculated by ASReml for each parameter are shown below each matrix. **G** matrices and standard errors are reprinted from McGlothlin et al. (2018a, b)). The full sampling (co)variance matrix, which was used to calculate the standard errors shown below, is available in our GitHub repository. Although we did not conduct formal likelihood-ratio tests, parameters that exceeded their standard errors by a factor of two are shown in bold, which provides a guide to statistical significance. Traits are abbreviated as follows: JL = jaw length, HW = head width, PECT = pectoral width, PELV = pelvic width, HUM = humerus, UL = ulna, FEM = femur, and TIB = tibia. All traits were natural-log transformed and size-corrected for analysis.

|  | JL | HW | PECT | PELV | HUM | UL | FEM | TIB |
| --- | --- | --- | --- | --- | --- | --- | --- | --- |
| JL | **.329** |  |  |  |  |  |  |  |
| HW | **.094** | **.449** |  |  |  |  |  |  |
| PECT | -.083 | **.349** | **1.426** |  |  |  |  |  |
| PELV | -.089 | **.240** | **.487** | **.546** |  |  |  |  |
| HUM | **.293** | .071 | **-.371** | **-.168** | **1.441** |  |  |  |
| UL | .079 | .075 | -.098 | .017 | **1.008** | **1.087** |  |  |
| FEM | **.208** | .030 | -.053 | -.051 | **.904** | **.731** | **.809** |  |
| TIB | **.268** | .009 | **-.172** | -.081 | **.945** | **.780** | **.783** | **.949** |

SE ( $\times 10^{-3}$ )
|  | JL | HW | PECT | PELV | HUM | UL | FEM | TIB |
| --- | --- | --- | --- | --- | --- | --- | --- | --- |
| JL | .052 |  |  |  |  |  |  |  |
| HW | .029 | .048 |  |  |  |  |  |  |
| PECT | .053 | .066 | .170 |  |  |  |  |  |
| PELV | .056 | .049 | .093 | .113 |  |  |  |  |
| HUM | .057 | .067 | .128 | .090 | .200 |  |  |  |
| UL | .053 | .063 | .119 | .087 | .144 | .175 |  |  |
| FEM | .035 | .042 | .077 | .058 | .093 | .085 | .071 |  |
| TIB | .037 | .043 | .081 | .060 | .096 | .089 | .067 | .076 |
| JL | **.458** |  |  |  |  |  |  |  |
| HW | **.245** | **.353** |  |  |  |  |  |  |
| PECT | -.122 | **.435** | **.306** |  |  |  |  |  |
| PELV | -.210 | **.484** | **.552** | **.196** |  |  |  |  |
| HUM | **.425** | .088 | **-.259** | -.190 | **.231** |  |  |  |
| UL | .132 | .108 | -.078 | .022 | **.805** | **.185** |  |  |
| FEM | **.403** | .049 | -.049 | -.076 | **.837** | **.779** | **.482** |  |
| TIB | **.480** | .014 | **-.148** | -.113 | **.808** | **.768** | **.894** | **.561** |

$h^2/r_g$ SE
|  | JL | HW | PECT | PELV | HUM | UL | FEM | TIB |
| --- | --- | --- | --- | --- | --- | --- | --- | --- |
| JL | .057 |  |  |  |  |  |  |  |
| HW | .068 | .028 |  |  |  |  |  |  |
| PECT | .079 | .073 | .029 |  |  |  |  |  |
| PELV | .130 | .093 | .092 | .037 |  |  |  |  |
| HUM | .081 | .082 | .084 | .101 | .028 |  |  |  |
| UL | .091 | .090 | .094 | .111 | .057 | .027 |  |  |
| FEM | .066 | .068 | .072 | .087 | .049 | .058 | .025 |  |
| TIB | .063 | .066 | .069 | .085 | .051 | .057 | .017 | .023 |

*Anolis evermanni* (Trunk-crown, Puerto Rico) $G (\times 10^{-3})$
|  | JL | HW | PECT | PELV | HUM | UL | FEM | TIB |
| --- | --- | --- | --- | --- | --- | --- | --- | --- |
| JL | **.251** |  |  |  |  |  |  |  |
| HW | -.013 | **.314** |  |  |  |  |  |  |
| PECT | .016 | **.411** | **1.062** |  |  |  |  |  |
| PELV | -.017 | **.183** | **.435** | **.623** |  |  |  |  |
| HUM | **.143** | .022 | .051 | .139 | **.866** |  |  |  |
| UL | **.141** | -.026 | .097 | .145 | **.884** | **.971** |  |  |
| FEM | .070 | .040 | **.228** | **.279** | **.850** | **.922** | **1.224** |  |
| TIB | **.121** | .047 | .178 | **.258** | **.971** | **1.064** | **1.427** | **1.751** |

$SE (\times 10^{-3})$
|  | JL | HW | PECT | PELV | HUM | UL | FEM | TIB |
| --- | --- | --- | --- | --- | --- | --- | --- | --- |
| JL | .029 |  |  |  |  |  |  |  |
| HW | .027 | .049 |  |  |  |  |  |  |
| PECT | .048 | .069 | .163 |  |  |  |  |  |
| PELV | .040 | .054 | .099 | .146 |  |  |  |  |
| HUM | .049 | .061 | .110 | .091 | .160 |  |  |  |
| UL | .049 | .062 | .113 | .093 | .135 | .156 |  |  |
| FEM | .041 | .054 | .098 | .083 | .108 | .110 | .117 |  |
| TIB | .048 | .064 | .114 | .097 | .124 | .128 | .125 | .174 |

$h^2/r_g$
|  | JL | HW | PECT | PELV | HUM | UL | FEM | TIB |
| --- | --- | --- | --- | --- | --- | --- | --- | --- |
| JL | **.415** |  |  |  |  |  |  |  |
| HW | -.047 | **.244** |  |  |  |  |  |  |
| PECT | .031 | **.712** | **.280** |  |  |  |  |  |
| PELV | -.043 | **.414** | **.535** | **.227** |  |  |  |  |
| HUM | **.306** | .042 | .053 | .189 | **.194** |  |  |  |
| UL | **.286** | -.046 | .096 | .186 | **.964** | **.212** |  |  |
| FEM | .126 | .064 | **.200** | **.319** | **.825** | **.846** | **.599** |  |
| TIB | **.182** | .064 | .131 | **.247** | **.788** | **.816** | **.975** | **.704** |

$h^2/r_g SE$
|  | JL | HW | PECT | PELV | HUM | UL | FEM | TIB |
| --- | --- | --- | --- | --- | --- | --- | --- | --- |
| JL | .033 |  |  |  |  |  |  |  |
| HW | .097 | .032 |  |  |  |  |  |  |
| PECT | .094 | .077 | .035 |  |  |  |  |  |
| PELV | .101 | .108 | .106 | .048 |  |  |  |  |
| HUM | .096 | .117 | .117 | .122 | .032 |  |  |  |
| UL | .092 | .113 | .111 | .117 | .044 | .029 |  |  |
| FEM | .072 | .087 | .083 | .089 | .066 | .053 | .030 |  |
| TIB | .069 | .085 | .083 | .089 | .071 | .057 | .011 | .034 |

*Anolis grahami* (Trunk-crown, Jamaica) $G (\times 10^{-3})$
|  | JL | HW | PECT | PELV | HUM | UL | FEM | TIB |
| --- | --- | --- | --- | --- | --- | --- | --- | --- |
| JL | **.246** |  |  |  |  |  |  |  |
| HW | **.218** | **.296** |  |  |  |  |  |  |
| PECT | .127 | .122 | **.351** |  |  |  |  |  |
| PELV | **.193** | .126 | .173 | **.704** |  |  |  |  |
| HUM | .152 | .067 | .167 | .264 | **1.003** |  |  |  |
| UL | .129 | .131 | .122 | .130 | **.856** | **1.171** |  |  |
| FEM | **.126** | -.005 | .081 | .186 | **.625** | **.559** | **.510** |  |
| TIB | **.146** | .034 | .171 | **.269** | **.732** | **.724** | **.549** | **.648** |

$SE (\times 10^{-3})$
|  | JL | HW | PECT | PELV | HUM | UL | FEM | TIB |
| --- | --- | --- | --- | --- | --- | --- | --- | --- |
| JL | .049 |  |  |  |  |  |  |  |
| HW | .051 | .082 |  |  |  |  |  |  |
| PECT | .064 | .081 | .151 |  |  |  |  |  |
| PELV | .068 | .084 | .113 | .169 |  |  |  |  |
| HUM | .140 | .186 | .238 | .176 | .465 |  |  |  |
| UL | .103 | .133 | .177 | .186 | .323 | .405 |  |  |
| FEM | .052 | .099 | .139 | .111 | .230 | .191 | .170 |  |
| TIB | .056 | .102 | .145 | .104 | .230 | .181 | .163 | .179 |

$h^2/r_g$
|  | JL | HW | PECT | PELV | HUM | UL | FEM | TIB |
| --- | --- | --- | --- | --- | --- | --- | --- | --- |
| JL | **.421** |  |  |  |  |  |  |  |
| HW | **.807** | **.240** |  |  |  |  |  |  |
| PECT | **.434** | .377 | **.133** |  |  |  |  |  |
| PELV | **.465** | .277 | .349 | **.277** |  |  |  |  |
| HUM | .307 | .123 | .281 | .315 | **.168** |  |  |  |
| UL | .240 | .222 | .191 | .143 | **.790** | **.159** |  |  |
| FEM | **.355** | -.013 | .191 | .311 | **.873** | **.723** | **.453** |  |
| TIB | **.367** | .076 | .358 | **.399** | **.908** | **.831** | **.955** | **.544** |

$h^2/r_g SE$
|  | JL | HW | PECT | PELV | HUM | UL | FEM | TIB |
| --- | --- | --- | --- | --- | --- | --- | --- | --- |
| JL | .056 |  |  |  |  |  |  |  |
| HW | .092 | .056 |  |  |  |  |  |  |
| PECT | .190 | .229 | .054 |  |  |  |  |  |
| PELV | .134 | .173 | .210 | .054 |  |  |  |  |
| HUM | .266 | .341 | .392 | .192 | .072 |  |  |  |
| UL | .188 | .221 | .281 | .199 | .155 | .050 |  |  |
| FEM | .136 | .249 | .323 | .171 | .165 | .160 | .122 |  |
| TIB | .127 | .231 | .289 | .134 | .165 | .124 | .037 | .110 |

*Anolis lineatopus* (Trunk-ground, Jamaica) $G (\times 10^{-3})$
|  | JL | HW | PECT | PELV | HUM | UL | FEM | TIB |
| --- | --- | --- | --- | --- | --- | --- | --- | --- |
| JL | **.442** |  |  |  |  |  |  |  |
| HW | **.286** | **.263** |  |  |  |  |  |  |
| PECT | **.385** | **.290** | **.750** |  |  |  |  |  |
| PELV | .193 | .132 | .343 | **.916** |  |  |  |  |
| HUM | **.321** | **.240** | .223 | -.016 | **.968** |  |  |  |
| UL | **.412** | **.297** | .308 | -.200 | **.942** | **1.047** |  |  |
| FEM | **.423** | .180 | **.547** | .228 | **.530** | **.540** | **1.013** |  |
| TIB | **.368** | .145 | **.528** | .182 | **.464** | **.495** | **.864** | **.825** |

$SE (\times 10^{-3})$
|  | JL | HW | PECT | PELV | HUM | UL | FEM | TIB |
| --- | --- | --- | --- | --- | --- | --- | --- | --- |
| JL | .103 |  |  |  |  |  |  |  |
| HW | .059 | .090 |  |  |  |  |  |  |
| PECT | .101 | .100 | .309 |  |  |  |  |  |
| PELV | .158 | .139 | .254 | .352 |  |  |  |  |
| HUM | .127 | .103 | .213 | .200 | .291 |  |  |  |
| UL | .106 | .104 | .186 | .174 | .240 | .270 |  |  |
| FEM | .168 | .130 | .233 | .256 | .255 | .248 | .346 |  |
| TIB | .147 | .117 | .212 | .227 | .226 | .222 | .295 | .272 |

$h^2/r_g$
|  | JL | HW | PECT | PELV | HUM | UL | FEM | TIB |
| --- | --- | --- | --- | --- | --- | --- | --- | --- |
| JL | **.536** |  |  |  |  |  |  |  |
| HW | **.837** | **.177** |  |  |  |  |  |  |
| PECT | **.668** | **.652** | **.151** |  |  |  |  |  |
| PELV | .303 | .269 | .414 | **.253** |  |  |  |  |
| HUM | **.491** | **.475** | .262 | -.017 | **.161** |  |  |  |
| UL | **.605** | **.566** | .348 | -.204 | **.936** | **.169** |  |  |
| FEM | **.633** | .349 | **.627** | .237 | **.535** | **.525** | **.645** |  |
| TIB | **.610** | .311 | **.672** | .209 | **.519** | **.533** | **.945** | **.587** |

$h^2/r_g SE$
|  | JL | HW | PECT | PELV | HUM | UL | FEM | TIB |
| --- | --- | --- | --- | --- | --- | --- | --- | --- |
| JL | .089 |  |  |  |  |  |  |  |
| HW | .114 | .056 |  |  |  |  |  |  |
| PECT | .158 | .206 | .059 |  |  |  |  |  |
| PELV | .217 | .265 | .263 | .087 |  |  |  |  |
| HUM | .167 | .196 | .251 | .223 | .044 |  |  |  |
| UL | .118 | .178 | .223 | .177 | .071 | .040 |  |  |
| FEM | .143 | .237 | .202 | .232 | .201 | .196 | .155 |  |
| TIB | .146 | .238 | .188 | .233 | .206 | .196 | .029 | .143 |

*Anolis pulchellus* (Grass-bush, Puerto Rico) $G (\times 10^{-3})$
|  | JL | HW | PECT | PELV | HUM | UL | FEM | TIB |
| --- | --- | --- | --- | --- | --- | --- | --- | --- |
| JL | .084 |  |  |  |  |  |  |  |
| HW | **.080** | **.157** |  |  |  |  |  |  |
| PECT | **.222** | **.215** | **.895** |  |  |  |  |  |
| PELV | .070 | .065 | .300 | **.766** |  |  |  |  |
| HUM | **.148** | .040 | .153 | -.036 | **.745** |  |  |  |
| UL | **.180** | .080 | .214 | -.152 | **.773** | **.982** |  |  |
| FEM | .092 | .040 | .156 | .126 | **.475** | **.524** | **.701** |  |
| TIB | .089 | .030 | .136 | .134 | **.403** | **.524** | **.648** | **.723** |

$SE (\times 10^{-3})$
|  | JL | HW | PECT | PELV | HUM | UL | FEM | TIB |
| --- | --- | --- | --- | --- | --- | --- | --- | --- |
| JL | .050 |  |  |  |  |  |  |  |
| HW | .040 | .059 |  |  |  |  |  |  |
| PECT | .075 | .070 | .268 |  |  |  |  |  |
| PELV | .066 | .061 | .192 | .244 |  |  |  |  |
| HUM | .072 | .068 | .163 | .127 | .204 |  |  |  |
| UL | .082 | .076 | .155 | .145 | .197 | .254 |  |  |
| FEM | .054 | .053 | .110 | .083 | .133 | .118 | .156 |  |
| TIB | .054 | .045 | .087 | .082 | .141 | .143 | .148 | .168 |

$h^2/r_g$
|  | JL | HW | PECT | PELV | HUM | UL | FEM | TIB |
| --- | --- | --- | --- | --- | --- | --- | --- | --- |
| JL | .076 |  |  |  |  |  |  |  |
| HW | **.701** | **.124** |  |  |  |  |  |  |
| PECT | **.810** | **.574** | **.211** |  |  |  |  |  |
| PELV | .276 | .187 | .363 | **.184** |  |  |  |  |
| HUM | .594 | .118 | .188 | -.047 | **.106** |  |  |  |
| UL | **.629** | .204 | .228 | -.175 | **.905** | **.116** |  |  |
| FEM | .382 | .122 | .196 | .172 | **.658** | **.632** | **.420** |  |
| TIB | .362 | .090 | .170 | .181 | **.549** | **.622** | **.911** | **.476** |

$h^2/r_g SE$
|  | JL | HW | PECT | PELV | HUM | UL | FEM | TIB |
| --- | --- | --- | --- | --- | --- | --- | --- | --- |
| JL | .044 |  |  |  |  |  |  |  |
| HW | .218 | .044 |  |  |  |  |  |  |
| PECT | .237 | .153 | .057 |  |  |  |  |  |
| PELV | .259 | .176 | .191 | .054 |  |  |  |  |
| HUM | .299 | .201 | .196 | .167 | .027 |  |  |  |
| UL | .295 | .192 | .164 | .160 | .066 | .028 |  |  |
| FEM | .211 | .158 | .134 | .113 | .127 | .113 | .074 |  |
| TIB | .214 | .134 | .108 | .112 | .159 | .130 | .038 | .086 |

*Anolis sagrei* (Trunk-ground, Bahamas) $G (\times 10^{-3})$
|  | JL | HW | PECT | PELV | HUM | UL | FEM | TIB |
| --- | --- | --- | --- | --- | --- | --- | --- | --- |
| JL | **.292** |  |  |  |  |  |  |  |
| HW | **.112** | **.302** |  |  |  |  |  |  |
| PECT | -.002 | **.166** | **.286** |  |  |  |  |  |
| PELV | -.077 | -.018 | .035 | **.463** |  |  |  |  |
| HUM | **.156** | -.073 | .027 | -.109 | **.603** |  |  |  |
| UL | .027 | **-.142** | -.047 | .082 | **.462** | **.646** |  |  |
| FEM | **.192** | -.033 | -.027 | -.011 | **.409** | **.315** | **.368** |  |
| TIB | **.177** | **-.111** | -.036 | -.003 | **.527** | **.507** | **.434** | **.606** |

$SE (\times 10^{-3})$
|  | JL | HW | PECT | PELV | HUM | UL | FEM | TIB |
| --- | --- | --- | --- | --- | --- | --- | --- | --- |
| JL | .023 |  |  |  |  |  |  |  |
| HW | .021 | .033 |  |  |  |  |  |  |
| PECT | .024 | .030 | .048 |  |  |  |  |  |
| PELV | .050 | .055 | .043 | .073 |  |  |  |  |
| HUM | .035 | .050 | .050 | .068 | .092 |  |  |  |
| UL | .037 | .044 | .053 | .062 | .080 | .116 |  |  |
| FEM | .024 | .037 | .042 | .041 | .051 | .051 | .054 |  |
| TIB | .026 | .044 | .051 | .042 | .058 | .061 | .056 | .073 |

$h^2/r_g$
|  | JL | HW | PECT | PELV | HUM | UL | FEM | TIB |
| --- | --- | --- | --- | --- | --- | --- | --- | --- |
| JL | **.469** |  |  |  |  |  |  |  |
| HW | **.377** | **.269** |  |  |  |  |  |  |
| PECT | -.007 | **.566** | **.127** |  |  |  |  |  |
| PELV | -.211 | -.049 | .095 | **.147** |  |  |  |  |
| HUM | **.372** | -.171 | .065 | -.206 | **.133** |  |  |  |
| UL | .063 | **-.321** | -.110 | .149 | **.740** | **.130** |  |  |
| FEM | **.585** | -.100 | -.085 | -.027 | **.869** | **.647** | **.270** |  |
| TIB | **.421** | **-.259** | -.087 | -.006 | **.873** | **.811** | **.919** | **.449** |

$h^2/r_g SE$
|  | JL | HW | PECT | PELV | HUM | UL | FEM | TIB |
| --- | --- | --- | --- | --- | --- | --- | --- | --- |
| JL | .023 |  |  |  |  |  |  |  |
| HW | .059 | .024 |  |  |  |  |  |  |
| PECT | .087 | .083 | .020 |  |  |  |  |  |
| PELV | .134 | .146 | .114 | .021 |  |  |  |  |
| HUM | .077 | .116 | .118 | .126 | .019 |  |  |  |
| UL | .084 | .097 | .123 | .112 | .069 | .022 |  |  |
| FEM | .059 | .112 | .131 | .097 | .055 | .083 | .034 |  |
| TIB | .054 | .097 | .122 | .084 | .048 | .060 | .026 | .041 |

*Anolis smaragdinus* (Trunk-crown, Bahamas) $G (\times 10^{-3})$
|  | JL | HW | PECT | PELV | HUM | UL | FEM | TIB |
| --- | --- | --- | --- | --- | --- | --- | --- | --- |
| JL | **.348** |  |  |  |  |  |  |  |
| HW | .054 | .161 |  |  |  |  |  |  |
| PECT | .251 | .065 | **.830** |  |  |  |  |  |
| PELV | .102 | -.002 | **.442** | **.560** |  |  |  |  |
| HUM | **.268** | .151 | .130 | .122 | .395 |  |  |  |
| UL | **.238** | .094 | .249 | .300 | .331 | .452 |  |  |
| FEM | .205 | .061 | .136 | .115 | .285 | .287 | **.344** |  |
| TIB | .296 | .008 | .221 | .278 | **.351** | **.423** | **.420** | **.609** |

$SE (\times 10^{-3})$
|  | JL | HW | PECT | PELV | HUM | UL | FEM | TIB |
| --- | --- | --- | --- | --- | --- | --- | --- | --- |
| JL | .129 |  |  |  |  |  |  |  |
| HW | .091 | .139 |  |  |  |  |  |  |
| PECT | .171 | .174 | .413 |  |  |  |  |  |
| PELV | .072 | .068 | .144 | .142 |  |  |  |  |
| HUM | .130 | .093 | .163 | .122 | .280 |  |  |  |
| UL | .098 | .134 | .176 | .175 | .181 | .286 |  |  |
| FEM | .114 | .085 | .184 | .127 | .149 | .149 | .165 |  |
| TIB | .125 | .072 | .205 | .141 | .120 | .132 | .158 | .197 |

$h^2/r_g$
|  | JL | HW | PECT | PELV | HUM | UL | FEM | TIB |
| --- | --- | --- | --- | --- | --- | --- | --- | --- |
| JL | **.455** |  |  |  |  |  |  |  |
| HW | .227 | .121 |  |  |  |  |  |  |
| PECT | .468 | .176 | **.254** |  |  |  |  |  |
| PELV | .230 | -.007 | **.648** | **.251** |  |  |  |  |
| HUM | **.722** | .600 | .228 | .259 | .100 |  |  |  |
| UL | **.601** | .348 | .406 | .596 | **.783** | .090 |  |  |
| FEM | **.592** | .257 | .255 | .262 | **.773** | **.728** | **.283** |  |
| TIB | **.644** | .025 | .311 | **.477** | **.717** | **.806** | **.919** | **.454** |

$h^2/r_g SE$
|  | JL | HW | PECT | PELV | HUM | UL | FEM | TIB |
| --- | --- | --- | --- | --- | --- | --- | --- | --- |
| JL | .137 |  |  |  |  |  |  |  |
| HW | .353 | .101 |  |  |  |  |  |  |
| PECT | .248 | .450 | .116 |  |  |  |  |  |
| PELV | .151 | .254 | .179 | .053 |  |  |  |  |
| HUM | .298 | .421 | .302 | .261 | .068 |  |  |  |
| UL | .235 | .481 | .296 | .306 | .290 | .055 |  |  |
| FEM | .207 | .348 | .337 | .274 | .307 | .311 | .122 |  |
| TIB | .154 | .223 | .279 | .205 | .269 | .243 | .094 | .118 |

**Table A2:**
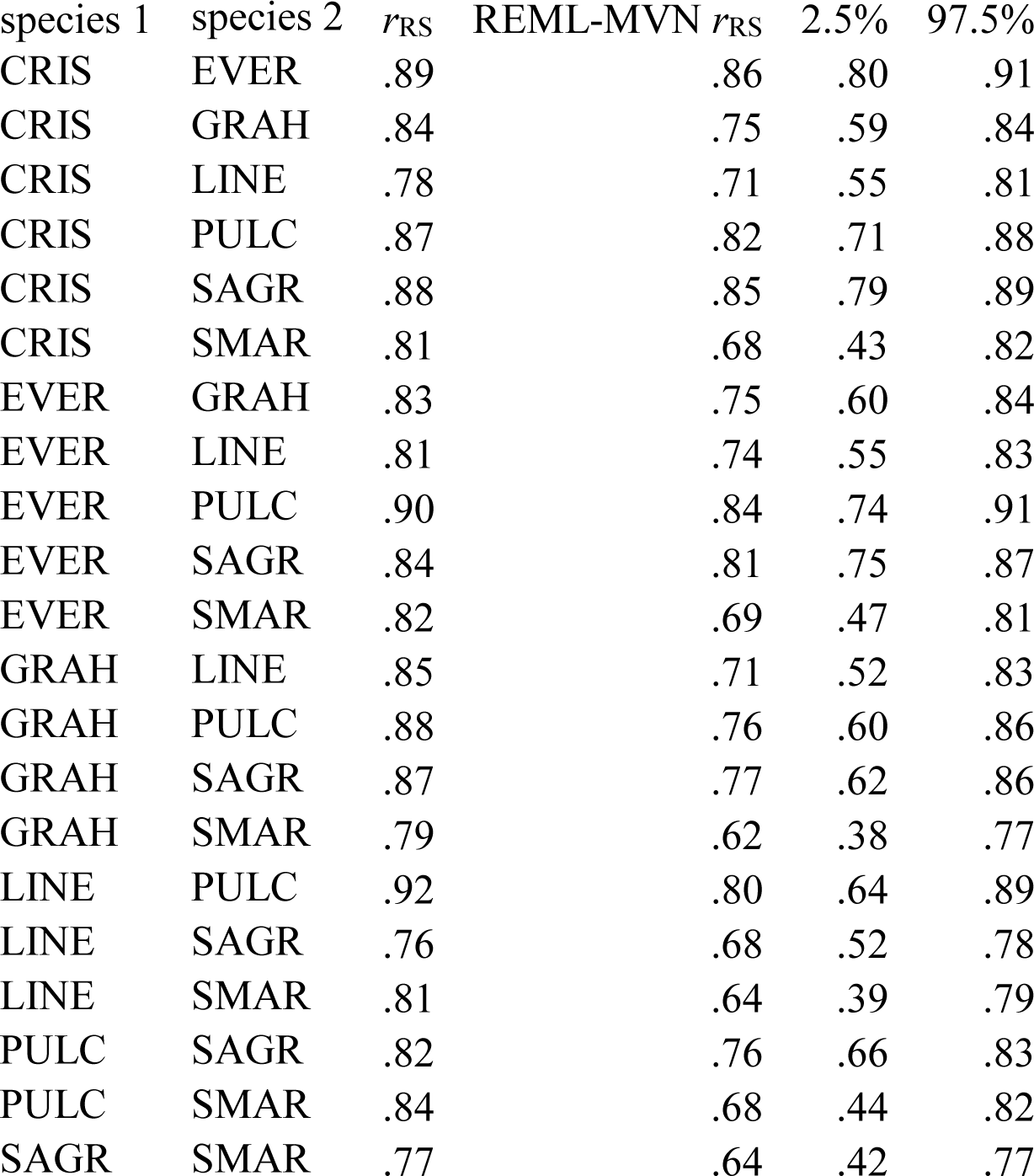
Random skewers correlations (*r*_RS_) for each species pair. Point estimates, REML-MVN estimates, and 95% confidence intervals are shown. Point estimates occasionally lie outside the REML-MVN confidence interval because incorporating estimation error from two **G** matrices lead to negatively skewed distributions.

**Table A3:** Estimates of phylogenetic signal (Blomberg’s *K*) for all genetic variances and correlations. Point estimates, REML-MVN estimates, and 95% confidence intervals are shown. All estimates of *K* were significantly greater than zero and statistically indistinguishable from 1, indicating phylogenetic signal consistent with a Brownian motion model of evolution.

Variances
| Trait | $K$ | REML-MVN $K$ | 2.5% | 97.5% |
| --- | --- | --- | --- | --- |
| JL | .81 | .85 | .65 | 1.11 |
| HW | .69 | .75 | .61 | 1.05 |
| PECT | 1.16 | 1.11 | .79 | 1.53 |
| PELV | .90 | .89 | .69 | 1.18 |
| RAD | .87 | .89 | .69 | 1.20 |
| UL | 1.15 | 1.03 | .77 | 1.31 |
| FEM | .99 | .98 | .83 | 1.31 |
| TIB | .90 | .90 | .77 | 1.13 |

Correlations
| Trait 1 | Trait 2 | $K$ | REML-MVN $K$ | 2.5% | 97.5% |
| --- | --- | --- | --- | --- | --- |
| JL | HW | .92 | .95 | .74 | 1.26 |
| JL | PECT | .71 | .73 | .64 | 1.00 |
| JL | PELV | .88 | .89 | .67 | 1.25 |
| JL | HUM | .91 | .91 | .69 | 1.20 |
| JL | UL | .78 | .81 | .67 | 1.12 |
| JL | FEM | 1.04 | .99 | .73 | 1.32 |
| JL | TIB | .95 | .91 | .73 | 1.17 |
| HW | PECT | .97 | .98 | .74 | 1.19 |
| HW | PELV | 1.04 | 1.00 | .71 | 1.39 |
| HW | HUM | 1.04 | 1.02 | .82 | 1.20 |
| HW | UL | .97 | .99 | .84 | 1.14 |
| HW | FEM | .91 | .96 | .78 | 1.20 |
| HW | TIB | .97 | .97 | .82 | 1.16 |
| PECT | PELV | 1.02 | .97 | .74 | 1.30 |
| PECT | HUM | .85 | .90 | .65 | 1.30 |
| PECT | UL | .92 | .94 | .71 | 1.19 |
| PECT | FEM | .91 | .95 | .77 | 1.18 |
| PECT | TIB | 1.02 | 1.01 | .80 | 1.24 |
| PELV | HUM | .90 | .91 | .75 | 1.15 |
| PELV | UL | 1.03 | 1.01 | .77 | 1.19 |
| PELV | FEM | .81 | .88 | .71 | 1.11 |
| PELV | TIB | .93 | .93 | .72 | 1.18 |
| HUM | UL | .90 | .99 | .76 | 1.22 |
| HUM | FEM | .78 | .90 | .72 | 1.17 |
| HUM | TIB | .76 | .85 | .71 | 1.13 |
| UL | FEM | .87 | .97 | .71 | 1.31 |
| UL | TIB | .80 | .89 | .71 | 1.17 |
| FEM | TIB | .79 | .92 | .72 | 1.14 |

**Table A4:** Ecomorph effects given as evolutionary correlations (*r*_e_) for all genetic variances and correlations. A positive value indicates that trunk-ground species have a higher value than trunk-crown species. Point estimates, REML-MVN estimates, and 95% confidence intervals are shown. Significant values are indicated with boldface.

| Trait | $r_e$ | REML-MVN $r_e$ | 2.5% | 97.5% |
| --- | --- | --- | --- | --- |
| JL | .62 | .54 | -.17 | .88 |
| HW | .52 | .43 | -.26 | .84 |
| PECT | .23 | .21 | -.27 | .56 |
| PELV | .03 | -.03 | -.85 | .58 |
| RAD | .44 | .39 | -.24 | .78 |
| UL | .09 | .08 | -.59 | .70 |
| FEM | .01 | .01 | -.49 | .34 |
| TIB | -.35 | -.35 | -.65 | .02 |

Correlations
| Trait 1 | Trait 2 | $r_e$ | REML-MVN $r_e$ | 2.5% | 97.5% |
| --- | --- | --- | --- | --- | --- |
| JL | HW | .28 | .26 | -.13 | .52 |
| JL | PECT | -.23 | -.22 | -.60 | .22 |
| JL | PELV | **-.56** | **-.52** | -.84 | -.03 |
| JL | HUM | .09 | .07 | -.64 | .79 |
| JL | UL | -.20 | -.19 | -.64 | .32 |
| JL | FEM | **.63** | **.58** | .05 | .90 |
| JL | TIB | .49 | .45 | -.06 | .84 |
| HW | PECT | .24 | .19 | -.73 | .57 |
| HW | PELV | .03 | .02 | -.60 | .58 |
| HW | HUM | -.13 | -.11 | -.63 | .52 |
| HW | UL | -.01 | .01 | -.47 | .50 |
| HW | FEM | .08 | .06 | -.68 | .64 |
| HW | TIB | -.07 | -.07 | -.75 | .48 |
| PECT | PELV | -.37 | -.30 | -.79 | .36 |
| PECT | HUM | -.55 | -.41 | -.86 | .39 |
| PECT | UL | -.44 | -.36 | -.85 | .35 |
| PECT | FEM | -.11 | -.09 | -.70 | .42 |
| PECT | TIB | -.24 | -.22 | -.68 | .28 |
| PELV | HUM | **-.96** | **-.83** | -.98 | -.32 |
| PELV | UL | **-.68** | **-.60** | -.89 | -.06 |
| PELV | FEM | **-.82** | **-.68** | -.97 | -.01 |
| PELV | TIB | **-.87** | **-.79** | -.97 | -.25 |
| HUM | UL | -.16 | -.12 | -.82 | .46 |
| HUM | FEM | -.37 | -.29 | -.72 | .59 |
| HUM | TIB | -.33 | -.27 | -.70 | .59 |
| UL | FEM | -.64 | -.42 | -.86 | .41 |
| UL | TIB | -.58 | -.44 | -.82 | .45 |
| FEM | TIB | -.65 | -.51 | -.94 | .20 |

